# Consequences of sustained modulation of VIP-expressing neurons in mouse corticolimbic regions on anxiety-related avoidance behaviors, social functioning, and associative fear learning

**DOI:** 10.64898/2026.09.30.755164

**Authors:** Marina Taylor, Minh Anh Nguyen, Diego Pascual Cuadrado, Yami George, Michael J. Schmeisser, Janina Hesse, Dilja Krueger-Burg

## Abstract

GABAergic inhibitory neurons play a central role in shaping network dynamics, and accordingly, their dysfunction is implicated in the pathophysiology of various psychiatric, neurodevelopmental, and neurological disorders. In the past decade, studies on vasoactive intestinal polypeptide-expressing inhibitory neurons (VIP^+^ neurons) have positioned them as critical components of this circuitry. In particular, VIP^+^ neurons have been proposed to function within a canonical disinhibitory circuit motif, preferentially inhibiting other interneuron subtypes, to ultimately modulate the gain and activity of local pyramidal neurons. However, their contribution to psychiatrically relevant behavioral endophenotypes is still not fully understood, limiting our ability to target these neurons for therapeutic purposes. In the present study, we systematically assessed the functional contribution of VIP^+^ neurons to affective and social behaviors. To this end, we used a chemogenetic approach to manipulate the activity of VIP^+^ neurons in core corticolimbic regions, including the medial prefrontal cortex (mPFC), basolateral amygdala (BLA), and ventral hippocampus (vHPC). The consequences of either activating or inhibiting VIP^+^ neurons in these regions were observed while mice underwent a battery of behavioral tasks assessing anxiety-related avoidance behaviors, social interest and recognition, and associative fear learning. While sustained modulation of VIP^+^ neurons had no significant effect on anxiety-related or social behaviors in any of the brain regions assessed, it produced a robust reduction in associative fear learning specifically in the mPFC. These findings demonstrate that the consequences of sustained VIP^+^ neuron modulation depend on both anatomical location and behavioural context. Moreover, they identify VIP^+^ neurons in the mPFC as potentially clinically relevant targets for interventions aimed at modulating fear-related circuits.

## Introduction

GABAergic inhibitory signaling plays a central role in shaping the flow of information throughout the brain, and accordingly, alterations in the function of GABAergic synapses and circuits have been linked to a wide range of psychiatric and neurodevelopmental disorders, including anxiety disorders, major depression, schizophrenia, and autism spectrum disorders [1–4]. Despite ample evidence for this link, however, surprisingly few therapeutic strategies are currently available that directly target or modulate the GABAergic system. Moreover, even those that do exist, such as the benzodiazepines, which act as positive allosteric modulators at GABA_A_ receptors, are known to result in severe side effects such as dependence, sedation, ataxia and fatigue [3, 5–7], substantially limiting their clinical applicability. These complications likely stem from the fact that the biological mechanisms underlying GABAergic system dysfunction in psychiatric and neurodevelopmental disorders are still insufficiently understood, highlighting the urgent need for additional research in this field.

A defining feature of the GABAergic system is its morphological and functional complexity. Numerous different GABAergic neuron subtypes exist, each of which contributes differently to the connectivity and computational output of local circuits and longe-range networks [8, 9]. Accordingly, elucidating the differential roles of these diverse inhibitory neuron subtypes in psychiatrically relevant behavioral circuits will be essential for identifying the mechanisms underlying psychiatric disorders. Moreover, given that many of the clinical limitations of current GABAergic pharmacotherapies likely result from their indiscriminate action at different subtypes of inhibitory neuron subtypes, a better understanding of these subtypes may also offer a unique opportunity for circuit-specific targeted therapeutic interventions with fewer debilitating side effects [5].

Within the corticolimbic circuits that have been particularly prominently linked to the etiology of psychiatric disorders [10, 11], GABAergic neurons are commonly divided into parvalbumin-expressing (PV^+^), somatostatin-expressing (SST^+^), and 5-HT3a receptor-expressing populations, the latter including vasoactive intestinal polypeptide-expressing (VIP^+^) neurons [8, 9]. While the contributions of corticolimbic PV^+^ and SST^+^ neurons to psychiatric endophenotypes have been extensively studied, substantially less is known in this regard with respect to the function of VIP^+^ neurons. Unlike most other cortical inhibitory interneuron subtypes, VIP^+^ neurons predominantly target other GABAergic neurons, placing them in a unique position to act as disinhibitory gates that exert powerful regulatory effects upon cortical network states [12–14]. In particular, it has been proposed that by inhibiting SST^+^ neurons, and to a lesser extent PV^+^ neurons, VIP^+^ neurons release pyramidal neurons from inhibition and increase the gain of relevant inputs. This mechanism has been reported to enhance performance on tasks requiring attention, sensory discrimination, and memory [14–18]. Accordingly, evidence indicates that VIP^+^ neuron dysfunction may contribute to behavioral phenotypes in animal models of neurodevelopmental and psychiatric disorders [19–26]. To date, however, a further investigation of this role is hampered by a lack of systematic studies identifying the behavioral circuits that are most vulnerable to VIP^+^ neuron dysfunction in a psychiatrically relevant context.

In the present study, we aimed to address this issue by providing a systematic and comparative overview of the behavioral consequences of sustained chemogenetic modulation of VIP^+^ neurons in mice in three core corticolimbic brain regions, i.e. the basolateral amygdala (BLA), medial prefrontal cortex (mPFC), and ventral hippocampus (vHPC). Within this context, we assessed multiple behavioral domains relevant to neurodevelopmental and psychiatric endophenotypes, including anxiety-related avoidance behaviors, social interest and recognition, and associative learning, using automated behavioral profiling to extend our analyses. Our findings provide an important basis for future studies to selectively assess VIP^+^ neuron dysfunction in animal models of neuropsychiatric disorders, as well as raising important considerations in light of recent efforts to develop AAV-mediated chemogenetic approaches as a direct therapeutic strategy in psychiatric and neurodevelopmental disorders [27–29].

## Methods

### Animals

VIP-IRES-Cre mice [30] were used to achieve Cre-dependent expression of viral DREADD constructs specifically in VIP^+^ neurons. Breeders were originally obtained from the Jackson Laboratory (JAX #010908) and were backcrossed onto a C57BL/6J background for at least six generations before experiments. Breeding was conducted at the Translational Animal Research Center (TARC), Johannes Gutenberg University Mainz. Animals were group-housed (2-5 mice per cage) and were kept on a 12 h light/dark cycle with food and water ad libitum. For experiments, VIP-Cre heterozygous male mice were bred from VIP-Cre heterozygous x VIP-Cre WT breeding pairs and were used for experiments between 8-15 weeks of age. For the social recognition task, male WT mice between 4–6 weeks of age from our breeding colony, i.e. also on a C57BL/6J background, were used as social stimulus mice. All behavioral experiments were conducted during the light cycle. The experimenter was blind to the experimental group throughout data acquisition and analysis. All procedures were approved by the Landesuntersuchungsamt Rheinland-Pfalz (TVA G 22-1-087) and were conducted in accordance with the guidelines of the welfare of experimental animal use issued by the federal government of Germany and the Johannes Gutenberg University Mainz.

### Viral vectors for DREADD constructs

Ready-to-use AAV5 particles were purchased from Addgene to express the following DREADD constructs in a Cre-dependent manner: pAAV-hSyn-DIO-mCherry control (#50459-AAV5), pAAV-hSyn-DIO-hM3D(Gq)-mCherry (#44361-AAV5), or pAAV-hSyn-DIO-hM4D(Gi)-mCherry (#44362-AAV5).

### Stereotactic surgery for AAV delivery

VIP-Cre mice were anesthetized using isoflurane (4.5% for induction, 1.5–2% for maintenance; Cp pharma, Catalog No. #1214, 1 ml/ml) in 1L/min O_2_. Injections of carprofen (Zoetis, Catalog No. #10000934, 10 mg/kg s.c. 30 min before surgery) and lidocaine (Bela-Pharm, Catalog No. #V134509, 5 mg/ml, 100 μl locally over the skull after incision) were delivered for analgesia. During surgery, eye ointment (Bepanthen, Bayer Vital GmbH, PZN: 01578675) was applied to avoid eye drying. Lidocaine (Aspen Pharmacare, PZN:13231379) was administered topically at the sites where the ear bars were positioned, and body temperature was maintained at 37 °C using a heating pad. For postoperative medication, carprofen (10 mg/kg in drinking water with 5% sucrose) was provided for three days. If necessary, mice received s.c. injections of 1% glucose in 0.9% saline, as well as carprofen according to the above dosage.

Mice were bilaterally injected with AAV particles using a stereotactic frame (Kopf Instruments Model 940). Injections were performed with a disposable glass pipette at a rate of 100 nl/min. The following coordinates relative to bregma were used: mPFC (AP: −2 mm, ML: ±0.2 mm, DV: 1.9 mm), BLA (AP: +1.6 mm, ML: ±3.3 mm, DV: 4.5 mm) and vHPC (AP: +3.15 mm, ML: ±3.3 mm, DV: 4.0 mm). The volume of injection was 300 nl (mPFC) or 200 nl (BLA and vHPC) per hemisphere. The pipette was left in place for 5 minutes after the injection to minimize backflow. After the pipette was removed, the skin was sutured, and animals were then placed in an incubation box at 37°C until they recovered their normal behaviour before being returned to their home cage. Behavioral testing took place three weeks after surgery to allow for recovery and viral expression.

### DCZ dilution and administration

Deschloroclozapine dihydrochloride (DCZ, HelloBio, catalog # HB9126) was used as an agonist for muscarinic-based DREADDs. A 2.5 mg/ml stock of DCZ in 0.9% saline was prepared before each battery of behavioral tasks and stored at −20 °C for a maximum of one month. Before behavioral experiments, a stock tube was brought to RT and diluted to a final working concentration of 0.01 mg/ml DCZ in 0.9% saline. Before each experiment, mice were single-housed for a minimum of 1 hour, and DCZ (100 μg/kg at 10 ml/kg, i.p.) was administered 30 min before each behavioral task.

### Behavioral paradigms

All animals underwent a standardized battery of behavioral tasks that included an elevated plus maze (EPM), an open field test (OFT), a social recognition task (SRT), and an auditory fear conditioning paradigm, each separated by 48-72 hours (see Figure 1a-d). Before each task, mice were single housed for at least 1 hour in a room next to the experimental room in order to avoid interference of the DCZ injections between animals. Animals were transferred between their home cage and the behavioral apparatus using a PVC tube to reduce stress during handling. After each behavioral task, animals were returned to their home cage simultaneously 30 min after the last cage mate had finished the experiment.

**Figure 1.**
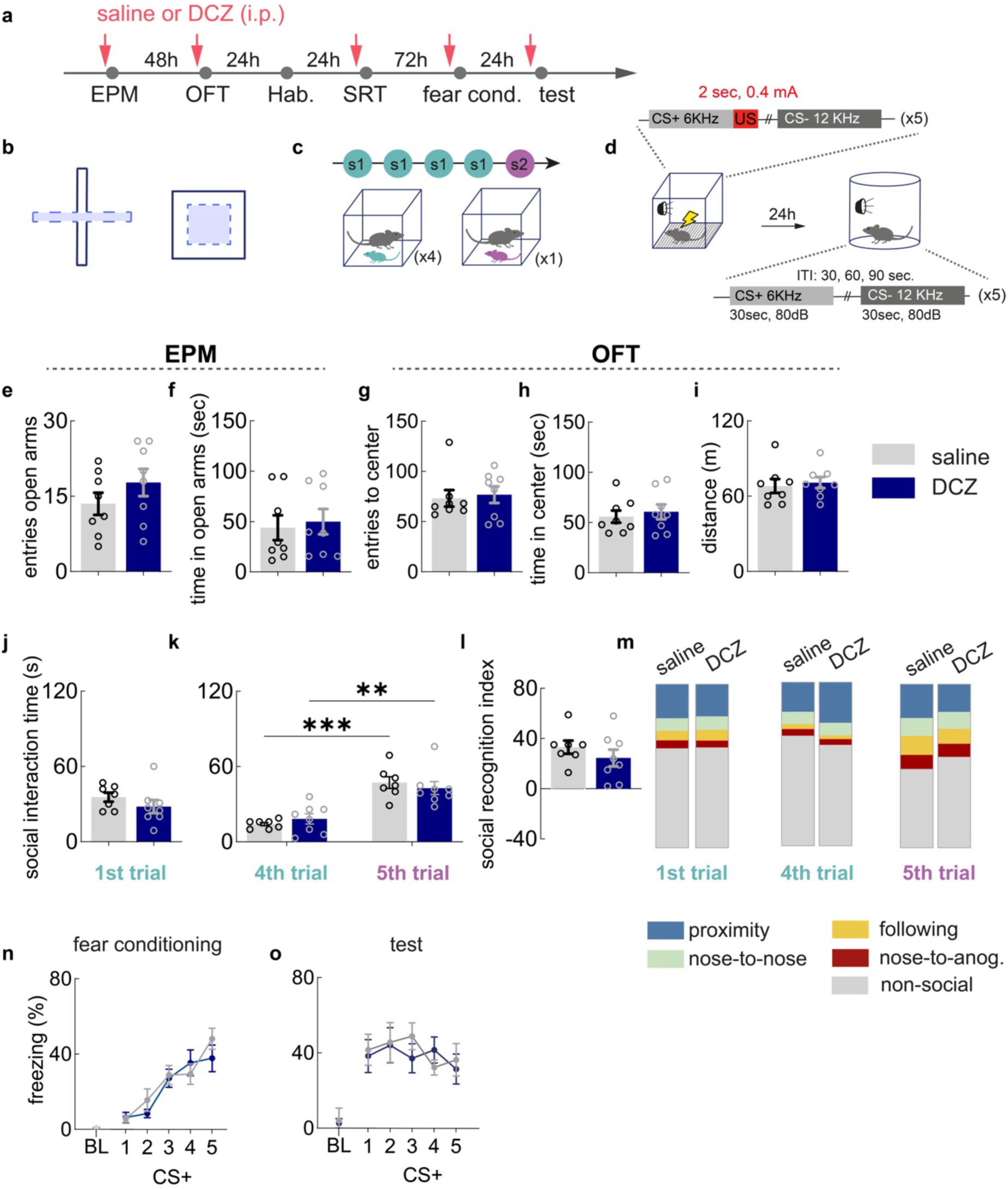
DCZ does not produce detectable off-target effects on behavioral measures in wild-type mice. **a.** Experimental timeline for saline or DCZ treatment and behavioral testing. **b**. Schematic of the elevated plus maze (EPM) and open field test (OFT) used to assess innate anxiety-like behavior. **c.** Schematic of the social recognition task (SRT), showing repeated exposure to the same stimulus mouse during Trials 1 to 4 and exposure to a novel stimulus mouse during Trial 5. **d.** Schematic of the fear-conditioning, and test protocol. **e.** Number of entries into the open arms of the EPM. **f.** Time spent in the open arms of the EPM, in seconds. **g.** Number of entries into the center of the OFT. **h.** Time spent in the center of the OFT, in seconds. **i.** Total distance traveled in the OFT, in meters. **j.** Time spent in social interaction during Trial 1 of SRT. **k.** Time spent in social interaction during Trials 4 and 5 of the SRT. **l.** social recognition index, calculated as social interaction time during Trial 5 minus social interaction time during Trial 4. **m.** video-based automated classification of mouse behavior during Trials 1, 4, and 5. Data are expressed as the percentage of the first 2 min of each trial spent exhibiting each behavior. **n.** Freezing during associative fear conditioning across CS+ time bins. **o.** Freezing during test session across CS+ time bins. Saline-treated mice are shown in light gray and DCZ-treated mice in dark blue. Data are presented as mean ± SEM. Statistically significant post-hoc comparisons are indicated by * p < 0.05, ** p < 0.01, *** p < 0.001, and **** p < 0.0001. Supplementary Table S1, S2 for statistical results.

#### Elevanted plus maze (EPM)

EPM testing was conducted in a white cross-shaped apparatus elevated 42 cm above the floor. The maze comprised two open arms (7 cm x 27 cm) and two closed arms (7 cm x 27 cm x 23 cm) arranged in opposite pairs, with a central intersection area (7 x 7 cm). At the beginning of the trial, mice were placed in the open arm facing the center of the apparatus, and they were allowed to investigate the apparatus for 5 minutes at a light intensity of 40 Lux. Performance was recorded using an overhead camera system (Basler ace GigE camera acA 1300-60gc), and time spent in the open arms was automatically scored using the EthoVision XT system (Noldus). Between mice, the apparatus was cleaned with water followed by 70% ethanol.

#### Open field test (OFT)

The OFT was performed in a square light grey plastic arena (50 cm x 50 cm x 40 cm) at a light intensity of 40 Lux. Mice were placed in the corner of the OF and allowed to investigate the arena for 10 minutes. Performance was recorded using an overhead camera system (Basler ace GigE camera acA 1300-60gc), and time spent in the center of the apparatus (defined as the central 30 cm x 30 cm area) as well as total distance travelled were automatically scored using the EthoVision XT system (Noldus). Between mice, the arena was cleaned with water followed by 70% ethanol.

#### Social Recognition Task (SRT)

The SRT, adapted from [31], was conducted in the same arena as the OFT at a light intensity of 40 Lux. 24 h before the SRT, experimental and stimulus mice were individually habituated to the arena for 10 min. The SRT itself consisted of 5 trials of 5 min each, separated by 5 min inter-trial intervals. In the first trial, a juvenile male mouse (stimulus mouse 1; 4-6 weeks old) was introduced to the experimental male mouse (12-14 weeks old), and they were allowed to interact freely. This was repeated for a total of 4 consecutive trials. In the 5th trial, a novel juvenile male mouse (stimulus mouse 2; 4-6 weeks old) was introduced to the experimental mouse. Performance was recorded using an overhead camera system. Social interaction was manually scored only for the first two minutes of the trial and only when the experimental mouse initiated the action. Aggressive behaviours were excluded. The arena was cleaned with water followed by 70% ethanol between trials and between mice. A Social Recognition Index (SRI) was calculated as (Trial 5 interaction time) -(Trial 4 interaction time).

#### Auditory fear conditioning paradigm

Auditory fear conditioning was performed using a Med Associates Mouse Fear Conditioning System (Med associates. Catalog no. 18372), which consisted of two fear conditioning chambers equipped with a near infrared video recording system. On day 1, mice were conditioned in context A, which consisted of a square chamber with a bar grid floor and metal walls (24 cm x 29 cm), placed into a white-walled sound attenuating chamber with a light intensity of 80 Lux. After a 2 min baseline phase, a pure tone (CS+: 6 kHz, 30 s, 80 dB) was delivered and paired with a US (2 s foot shock, 0.4 mA). Another pure tone (CS-: 12 kHz, 30 s, 80 dB) was presented interleaved without shock. The CS+/US and CS- were alternately presented five times with a pseudorandom inter-trial interval (30, 60, or 90 s), starting with the CS+. Between mice, the chamber was cleaned with water and 1 % acetic acid. On day 2, expression of the conditioned fear memory was tested. To this end, mice were placed in context B, which consisted of a semicircular chamber with a smooth white plastic floor and walls (24 cm x 29 cm), placed into a white-walled sound attenuating chamber with a light intensity of 20 Lux. After a 2 min baseline phase, the CS+ and CS- were alternately presented five times, but no US was delivered. The cage was cleaned with water and 70 % ethanol between mice. On both days, animals remained in the context for 1 min after the last CS presentation. Performance was recorded using the Med Associates VFC system, and freezing was automatically scored by Med Associates software. Freezing was detected using the Linear method and was counted when the animal was immobile for longer than 0.7 sec.

#### Automated behavioural profiling

Objective quantification of mouse behavior in the SRT was performed using video-based computational analysis (see Supplementary Methods for details). Body part tracking was conducted with DeepLabCut [32, 33], trained for the task and with additional post-processing by custom-made python scripts to improve mouse identity assignment, interpolate untracked body part positions, and minimize frame-to-frame tracking noise (see also Supplementary Figure S3). The behavioural classification was quantified by custom Python-based algorithms. For the SRT, interactions between mice were classified as *following* when the experimental mouse entered a trajectory ROI laid by the stimulus mouse, as *nose-to-nose* and *nose-to-tail* when the nose of the experimental mouse was in proximity of the nose or tailbase of the social stimulus mouse, and otherwise as *proximity* when the mice interacted.

### Immunohistochemistry

Following completion of the behavioral testing battery, animals were sacrificed for histological validation of virus placement. In order to additionally validate functionality of DREADD constructs using cFos immunostaining (see below), mice received an additional i.p. DCZ injection 90 min prior to sacrifice. Anesthesia was induced using ketamine (200 mg/kg) / xylazine (20 mg/kg) injected i.p. at 10 ml/kg. Mice were then perfused transcardially with 4% paraformaldehyde (PFA, ROTI®Histofix, ROTH, Catalog #P087.1) in PBS. Brains were extracted and post-fixed in 4% PFA for 24 h and transferred to 1x PBS containing 0.01% sodium azide for storage. Before cryosectioning, brains were transferred to 30% sucrose and allowed to equilibrate at 4°C. Subsequently, 40 μm thick tissue sections containing mPFC, BLA or vHPC were collected using a cryostat (Leica CM1950).

#### Immunohistochemistry for virus placement and cFos analysis

Sections were washed 3x 10 min with 1x PBS at RT, and incubated for 1.5 hours in blocking solution (1x PBS, 10% NGS, 0.3% v/v Triton X-100, 3% w/v BSA) at RT. Sections were incubated overnight at 4°C with primary antibodies (anti-cFos, guinea pig, 1:1000, 0.5 mg/ml, Synaptic Systems, catalog #226308; and anti-RFP, rabbit, 1:2000, 1 mg/ml, Rockland Immunochemicals, catalog # 600- 401-379) diluted in incubation solution (1x PBS, 5% NGS, 0.3% v/v Tritón X-100, 3% w/v BSA). The next day, sections were washed 3x 10 min with 1x PBS and incubated for 2 hours with secondary antibodies (anti-guinea pig antibody labelled with Alexa 488, 1:1200, 2 mg/ml, catalog #A11073 Invitrogen; and anti-rabbit antibody labelled with Alexa 647, 1:1200, 2 mg/ml, catalog #A32733 Invitrogen) diluted in incubation solution. Sections were washed, incubated for 10 min with DAPI (1:1000, 1 mg/ml, catalog #10374168, Thermofisher), washed again, and mounted on microscope slides with mounting media ROTI Mount FluorCare (Carl Roth, No. HP19.1) and coverslipped. Each microscope slide contained several brain sections from the targeted area of injection, as well as anterior and posterior regions to analyze the viral dispersion. Sections were kept at 4°C until imaging.

#### Immunohistochemistry for colocalization of VIP and mCherry

Sections were incubated in 10 mM citrate buffer containing 0.05% Triton X-100, pH 6.0, for 30 min at 95°C. They were then washed 3x 10 min with 1x PBS at RT and incubated for 2 h in blocking solution (1x PBS, 10% NGS, 0.3% v/v Tritón X-100) at RT. Sections were incubated for 72 hours at 4°C in primary antibodies (anti-RFP, chicken, 1:1000, 1 mg/ml, Rockland catalog #600-901-379; and anti-VIP, rabbit, 1:1000, 1 mg/ml, Immunostar catalog #20077) diluted in blocking solution. Sections were washed 3x with 0.1% Triton X-100 in 1x PBS at RT, and they were subsequently incubated with secondary antibodies (Alexa 488, anti-rabbit, 1:750, 2 mg/ml, catalog # A- 11008 Thermofisher; and Alexa 647, anti-Chicken, 1:750, 2 mg/ml, catalog # A-21449 Thermofisher) 24 hours at 4°C. Sections were then washed and incubated for 10 min with DAPI (1:1000, 1 mg/ml, catalog # 10374168, Thermofisher), washed again, and mounted as previously described.

### Image acquisition and analysis

The mounted tissue sections described above were imaged using a Leica SP8 confocal microscope (Leica Microsystems, Germany), equipped with both a hybrid detector (HyD) and a photomultiplier tube (PMT) for signal detection. The imaging parameters were consistent within one imaging session and between experimental groups. Analysis was performed using Fiji Image Software.

#### Image acquisition and analysis for assessment of virus placement

A tile scan of the slide containing at least 6 coronal brain sections was performed. Single plane images were acquired using a 10X objective with a 1X zoom and 256 × 256 pixel spatial resolution. DAPI was used to visualize overall brain anatomy, while the mCherry signal, amplified using an anti-RFP antibody, indicated the viral expression pattern. Animals were excluded from behavioral statistics if they did not meet the following criteria: bilateral virus expression, and target coverage of at least the prelimbic area (mPFC), basal amygdala (BA), or ventral CA1 (vHPC).

For animals with confirmed expression, expression site centers were plotted onto a coronal template of the target region (Supplementary Figure S1).

#### Image acquisition and analysis of cFos expression in DREADD-expressing neurons

Single plane images were taken using a 20X oil immersion objective with a 1X zoom and 512 x 512 spatial resolution. DAPI was used to visualize overall brain anatomy, while the mCherry signal indicated neurons expressing DREADD, and cFos indicated neuronal activation levels. Cells were classified as cFos^+^ at ≥ 7× background intensity, which was calculated for each animal from two random background areas per image. Images were binarized and despeckled, and cFos^+^ cells were counted with the function “Analyze Particles” using size (20–120 µm²) and circularity (0.30–1.00) thresholds. mCherry^+^ cells were counted manually, and a cFos^+^ mask was overlaid on mCherry images to identify double-positive (mCherry^+^/cFos^+^) cells.

#### Image acquisition and analysis for colocalization of VIP and mCherry immunostaining

Images were taken using a 63X oil immersion objective with a 1X zoom and 1024 x 1024 spatial resolution. A 10 µm Z-stack of 7 steps was performed in every tile. DAPI was used to visualize cellular nuclei, mCherry indicated neurons expressing DREADD, and VIP indicated VIP^+^ neurons. To assess mCherry specificity for VIP^+^ neurons, colocalization was analyzed on maximal z-stack projections. mCherry^+^ cells were manually counted (“Cell Count” tool), and the overlay with the VIP signal identified double-positive (mCherry^+^/VIP^+^) cells.

### Statistical analysis

Data sets were tested for normality. Outliers were identified using the Grubbs test, and animals were excluded if flagged as outliers for at least one variable in two or more behavioral experiments. Independent data sets were analyzed by one-way ANOVA (parametric) or Kruskal-Wallis (non-parametric), with Tukey’s or Dunn’s post hoc tests, respectively, following significant main effects. Dependent variables were analyzed by two-way repeated measures ANOVA, with Tukey’s post hoc test following significant main effects or interactions. Analyses were performed in GraphPad Prism 10.0. Graphs show means ± SEM, with significance set at p ≤ 0.05. The same approach was applied for the analysis of DCZ-driven DREADD activation and associated cFos expression, and for the effects of VIP^+^ neuron modulation on behavior (one-way ANOVA/Kruskal-Wallis for single independent variables across the three groups; two-way repeated measures ANOVA for two independent factors).

## Results

### Confirmation of lack of off-target effects of DCZ on behavior

In the current study, we investigated VIP^+^ neuron function in psychiatrically relevant behaviors using a chemogenetic approach based on activation or inhibition of VIP^+^ neurons through virally expressed Designer Receptors Exclusively Activated by Designer Drugs (DREADDs). A common concern with the use of this methodology is that many DREADD ligands, including the frequently used clozapine-N-oxide (CNO), are metabolized *in vivo* to drugs with pharmacological activity at endogenous receptors, resulting in off-target effects of the DREADD ligand itself on behavior [34]. To minimize these potential confounding effects, we used deschloroclozapine (DCZ), a next-generation DREADD ligand with greater potency and selectivity for DREADD receptors than CNO that has been reported to produce minimal off-target behavioral effects in rodents and non-human primates [35–37]. To confirm that DCZ administration alone does not affect the behavioral paradigms under investigation in the present study, we first assessed its effects on the relevant behavioral outcome measures in WT mice. To this end, WT male on a C57BL/6J background were tested on a behavioral battery (Figure 1a) including anxiety-related behaviors (elevated plus maze EPM, and open-field test, OFT, Figure 1b), social behavior (social recognition task, SRT, Figure 1c), and associative fear learning (auditory fear conditioning paradigm, Figure 1d). 30 min before each behavioral task, they received intraperitoneal injections of either DCZ (100 μg/kg in 0.9% saline) or saline vehicle. DCZ administration did not significantly affect the number of entries into or the time spent in the open arms of the EPM (Figure 1e, f), the number of entries into the center zone, the time spent in the center zone, or the distance traveled in the OFT (Figure 1g-i), indicating that DCZ administration did not affect anxiety-like or locomotor behaviour in WT mice. Similarly, DCZ administration did not affect social interest (Figure 1j), social recognition (Figure 1k, l), or the distribution of specific behavioral elements identified by a DeepLabCut-based behavioral analysis pipeline (Figure 1m) in WT mice. During associative fear learning, DCZ administration did not significantly affect freezing during fear acquisition (Figure 1n) or memory retrieval (Figure 1o). Together, these findings suggest that, at the dose and timing used, DCZ did not produce detectable off-target effects on the behavioral measures assessed in wild-type mice. These control data support the interpretation that behavioral effects observed in DREADD-expressing animals reflect receptor-mediated neuronal modulation rather than nonspecific effects of DCZ.

### Validation of DCZ-induced activation of DREADD-expressing VIP^+^ neurons in the BLA, mPFC and vHPC

A key objective of our study was to compare the behavioral function of VIP^+^ neurons in three different regions of the corticolimbic circuitry, i.e. the BLA, the mPFC, and the vHPC. To this end, we stereotaxically injected Cre-dependent DREADD-expressing AAV constructs into VIP-Cre mice (Figure 2a). Three AAV constructs were injected (Figure 2b): (1) a soluble mCherry control lacking a DREADD receptor and therefore insensitive to DCZ; (2) hM3Dq, a Gq-coupled muscarinic DREADD that increases the excitability of receptor-expressing neurons following ligand administration; and (3) hM4Di, a Gi-coupled muscarinic DREADD that decreases the excitability of receptor-expressing neurons following ligand administration [38]. Each of the three AAV constructs was injected into three independent cohorts of mice (one each for the BLA, the mPFC, and the vHPC), yielding a total of 9 experimental groups (Figure 2b-g). After three weeks for recovery and viral expression, mice were tested on the same behavioral test battery as described above (see also Figure 1a, results described below), and they were then sacrificed for analysis of viral placement (Supplementary Figure S1). Only mice that showed correct virus expression were included in the analysis of the following behavioral data.

**Figure 2.**
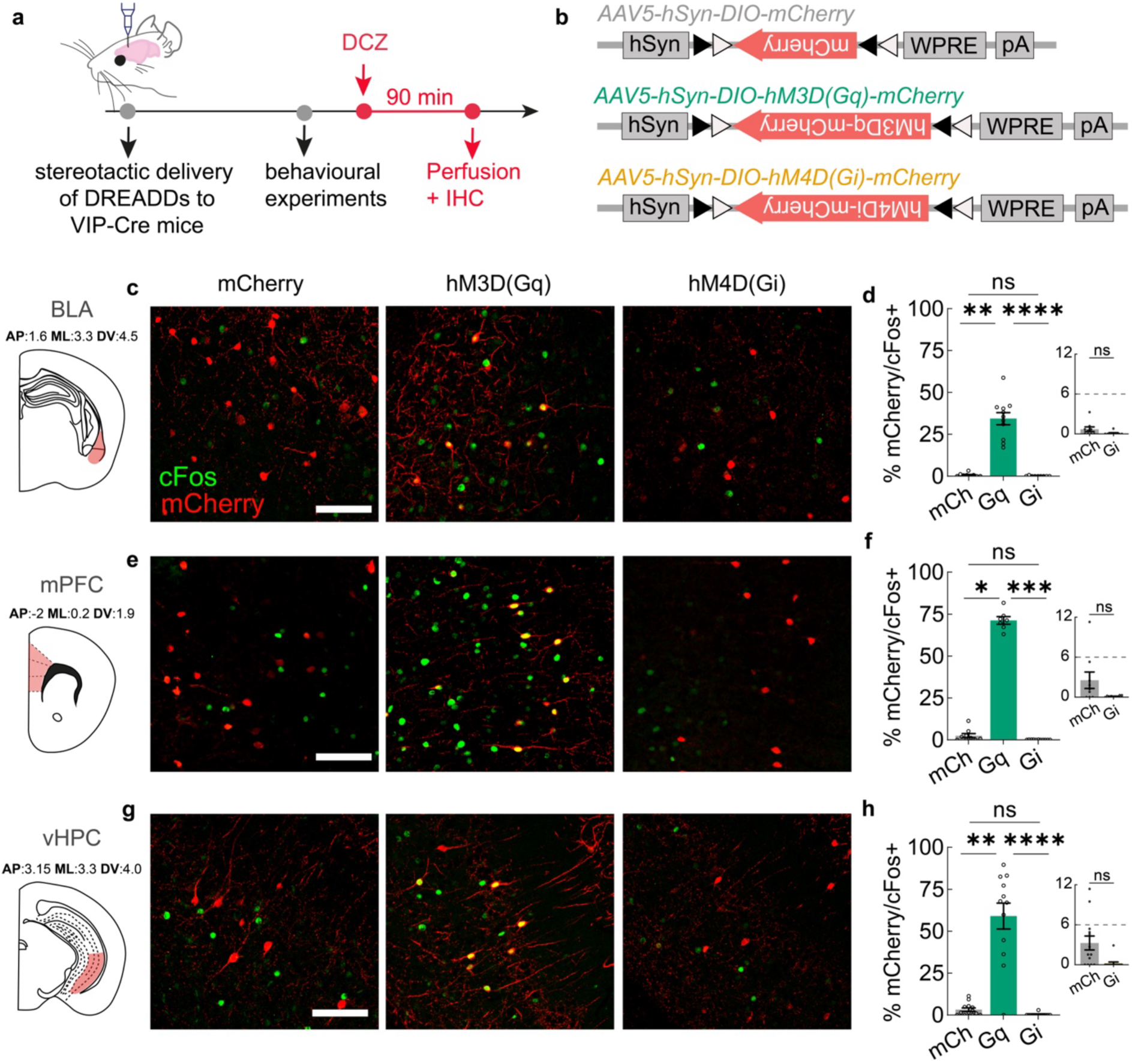
Validation of DCZ-induced activation of DREADD-expressing VIP neurons in the BLA, mPFC and vHPC. **a.** Experimental timeline showing stereotactic DREADD delivery to the basolateral amygdala (BLA), medial prefrontal cortex (mPFC), or ventral hippocampus (vHPC), behavioral testing, DCZ administration, perfusion 90 min later, and immunohistochemical detection of mCherry and cFos. **b.** Schematic of the viral construct used for targeted expression. c.Representative confocal images showing mCherry expression (red), cFos immunoreactivity (green), and colocalization (yellow) in the BLA across the mCherry-control, hM3D(Gq), and hM4D(Gi) groups. **d.** Percentage of mCherry-positive cells that were also cFos-positive in the BLA, quantified within an area of 0.20 mm². **e.** Representative confocal images showing mCherry expression (red), cFos immunoreactivity (green), and colocalization (yellow) in the mPFC across experimental groups. **f.** Percentage of mCherry-positive cells that were also cFos-positive in the mPFC, quantified within an area of 0.34 mm². **g.** Representative confocal images showing mCherry expression (red), cFos immunoreactivity (green), and colocalization (yellow) in the vHPC across experimental groups. **h.** Percentage of mCherry-positive cells that were also cFos-positive in the vHPC. BLA, basolateral amygdala; mPFC, medial prefrontal cortex; vHPC, ventral hippocampus. Scale bar, 100 µm. Data are presented as mean ± SEM. Statistically significant post hoc comparisons are indicated by * p < 0.05, ** p < 0.01, *** p < 0.001, and **** p < 0.0001. Supplementary Table S3 for statistical results.

To validate efficiency of viral transduction, the total number of mCherry^+^ cells was quantified (Supplementary Figure S2a, c, e,). In all three brain regions, the density of mCherry^+^ cells was similar across AAV constructs, although the hM3D(Gq) group exhibited a slightly lower density of mCherry^+^ cells than the control group in the mPFC and the vHPC (Supplementary Figure S2a, c, e). These findings show that transduction efficiency was similar, albeit not identical, across AAV constructs. To assess the specificity of DREADD expression, sections were immunostained for VIP and mCherry. Quantification of double-labeled cells revealed that DREADD expression was almost exclusively restricted to VIP-immunopositive neurons (Supplementary Figure S2g, h), confirming specificity of our experimental approach. To validate the DCZ-induced activation or inhibition of DREADD-expressing neurons, we additionally performed a neuronal activation analysis using cFos expression as a marker of neuronal activation. To this end, mice were injected with saline or DCZ 90 min prior to sacrifice, and immunohistochemical colocalization analysis of cFos with mCherry was used to assess the activation or inhibition of DREADD-expressing neurons following intraperitoneal DCZ administration (Figure 2c-h, Supplementary Figure S2). In all the brain regions, DCZ administration resulted in a pronounced increase in the proportion of mCherry^+^/cFos^+^ cells in the hM3D(Gq) group compared to both the control and hM4D(Gi) groups (BLA, Figure 2c, d; mPFC, Figure 2e, f; vHPC, Figure 2g, h). The number of mCherry^+^/cFos^+^ in the hM3D(Gq) group was largest in the mPFC (71% of mCherry^+^ neurons), followed. by vHPC (59% of mCherry^+^ neurons) and BLA (34% of mCherry^+^ neurons). Conversely, virtually no mCherry^+^/cFos^+^ cells were observed in the hM4D(Gi) group in all three brain regions (Figure 2c-h), indicating effective inhibition of neuronal activity following DCZ administration in this group. While the reduction of mCherry^+^/cFos^+^ cells in the hM4D(Gi) group compared to the control group did not reach statistical significance, it should be noted that the proportion of mCherry^+^/cFos^+^ cells was already very low in the control group across all three brain regions. This observation indicated that activity in the targeted VIP^+^ neurons was low under the baseline conditions assessed in the present analysis, which likely precluded the assessment of a further significant reduction in the hM4D(Gi) group.

Interestingly, the total number of cFos^+^ neurons was also increased in all three brain regions in the hM3D(Gq) group compared to the control and hM4D(Gi) groups (Supplementary Figure S2b, d, f), presumably reflecting the proposed disinhibitory function of VIP^+^ neurons upon activation. Strikingly, this increase in cFos^+^ neurons in the hM3D(Gq) group was substantially higher in the mPFC than in the BLA or the vHPC, potentially indicating that activation of VIP^+^ neurons exerts a more powerful influence over local network activity in the mPFC than in the BLA or the vHPC. No differences were observed between the control and hM4D(Gi) groups, consistent with the notion that under the baseline conditions of this experiment, insufficient VIP^+^ neurons are active under control conditions to be able to observe a further reduction due to inhibition of VIP^+^ neurons in the hM4D(Gi) group.

Together, these data validate the specificity and efficacy of our experimental approach and indicate that DCZ administration induces effective modulation of DREADD-expressing VIP^+^ neurons across the BLA, mPFC, and vHPC.

### Chemogenetic modulation of VIP^+^ neurons does not significantly alter anxiety-related behaviors

To examine whether VIP^+^ neurons contribute to the modulation of innate anxiety-related behaviors, performance was assessed in two tasks related to innate avoidance behaviors, the EPM and the OFT. To this end, DCZ was administered to control, hM3D(Gq) and hM4D(Gi) groups as previously validated, and 30 min later, mice were tested in either the EPM or the OFT (Figure 3a). In the EPM, anxiety-related behavior can be inferred from the innate avoidance of the open, anxiogenic arms. Neither activation nor inhibition of VIP^+^ neurons in any of the three brain regions significantly altered the time spent in the open arms (Figure 3b-d). Similarly, increased thigmotaxis in the OFT can be interpreted as anxiety-related behavior, since rodents preferentially avoid the anxiogenic open center of the arena. Again, modulation of VIP^+^ neurons in any of the three brain regions did not significantly affect the time spent in the center of the arena (Figure 3e-g). Moreover, no significant differences in locomotor activity were observed among groups (Figure 3h-j). Together, these results indicate that sustained chemogenetic modulation of VIP^+^ neurons in the BLA, mPFC or vHPC did not detectably alter anxiety-related avoidance behavior or locomotion under the experimental conditions tested.

**Figure 3.**
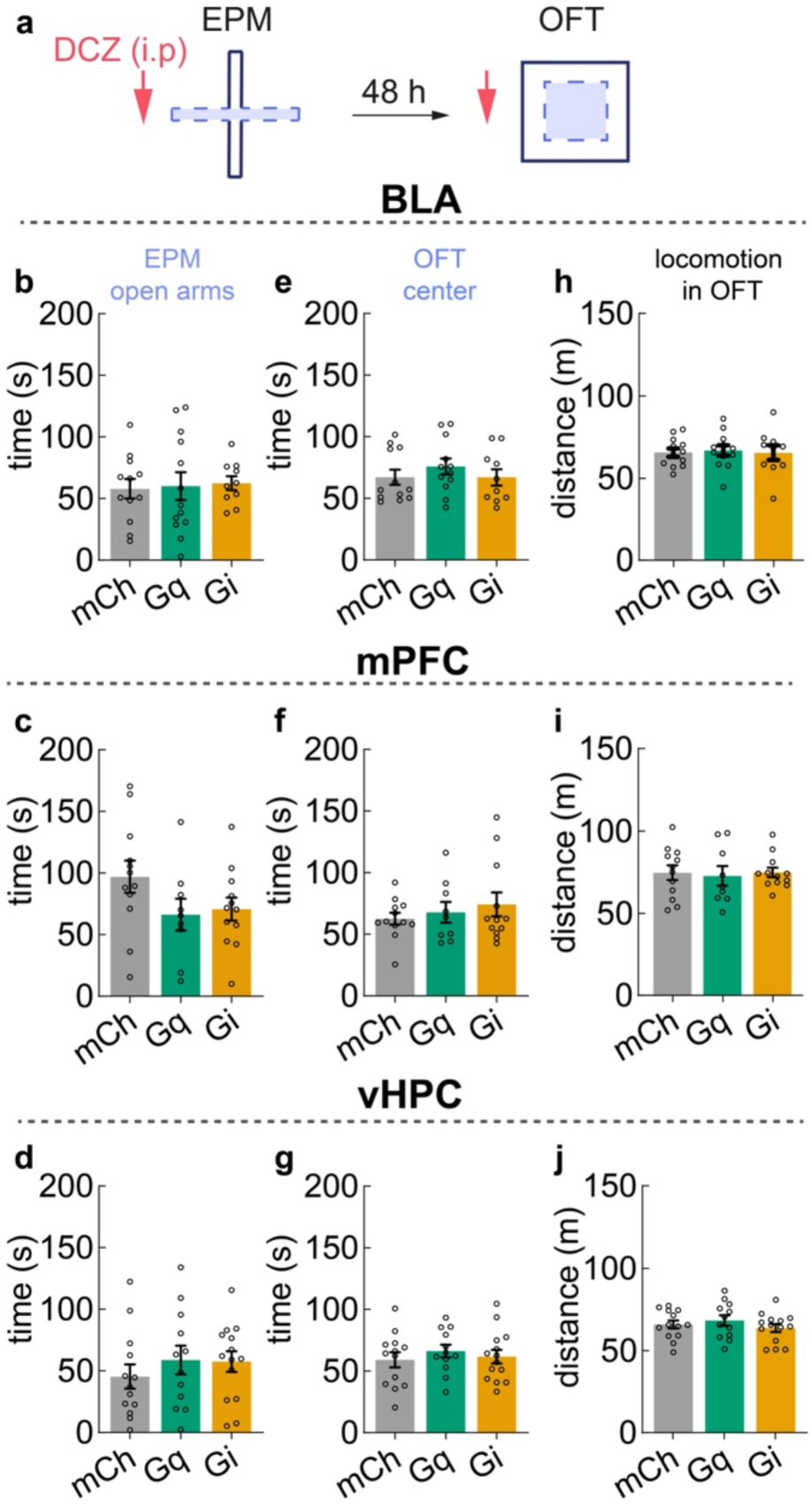
Sustained manipulation of VIP-expressing neurons does not significantly alter anxiety-related behavioral measures. **a.** Experimental timeline showing intraperitoneal DCZ administration, testing in the EPM, and OFT 48 h later. **b.** Time spent in the open arms of the EPM following manipulation of VIP-expressing neurons in the BLA. **c.** Time spent in the open arms of the EPM following manipulation in the mPFC. **d.** Time spent in the open arms of the EPM following manipulation in the vHPC. **e.** Time spent in the center of the OFT following manipulation in the BLA. **f.** Time spent in the center of the OFT following manipulation in the mPFC. **g.** Time spent in the center of the OFT following manipulation in the vHPC. **h.** Total distance traveled in the OFT following manipulation in the BLA. **i.** Total distance traveled in the OFT following manipulation in the mPFC. **j.** Total distance traveled in the OFT following manipulation in the vHPC. Data are presented as mean ± SEM. mCherry, hM3D(Gq), and hM4D(Gi) indicate the respective experimental groups. BLA, basolateral amygdala; mPFC, medial prefrontal cortex; EPM, elevated plus maze; OFT, open-field test. Supplementary Table S4 for statistical results.

### Chemogenetic modulation of VIP^+^ neurons does not significantly alter social interest or social recognition

Social impairment is a core symptom and often one of the earliest signs of psychiatric disorders, including schizophrenia and major depressive disorder [39], yet the neural mechanisms underlying social deficits remain poorly understood. In particular, almost nothing is known to date about the role of VIP^+^ neurons in social behavior. To begin to address this issue, we investigated the contribution of VIP^+^ neurons to social functioning using a social recognition task (Figure 4a, adapted from [31]). In this task, experimental mice were exposed to an unfamiliar juvenile mouse (stimulus mouse 1) in an open field arena for 4x 5 min trials, separated by 5 min inter-trial intervals, and interaction time initiated by the experimental mouse was assessed during the first two minutes of the first and last trial (i.e. trials 1 and 4). Following another 5 min inter-trial interval, experimental mice were exposed to a second unfamiliar juvenile mouse (stimulus mouse 2) for 5 min, and interaction time initiated by the experimental mouse was again assessed during the first two minutes (trial 5, Figure 4a). This task allowed us to assess two dimensions of mouse social behavior, i.e. social interest (defined as interaction time in trial 1) and social recognition (defined as the difference in interaction time between trial 4 and trial 5), in freely moving animals. To determine whether VIP^+^ neurons in the BLA, mPFC or vHPC contribute to either social interest or recognition in the SRT, VIP^+^ neurons were activated or inhibited chemogenetically compared with control animals during task performance as described above. Social behaviors were analyzed during the first two minutes of the trial, when the animals engaged in the highest level of social interaction (Supplementary Figure S4 a-j). Chemogenetic modulation of VIP^+^ neurons in the BLA, mPFC or vHPC did not significantly affect the time spent interacting socially during trial 1, indicating that baseline interest in social interaction was comparable across groups (Figure 4b-d). Moreover, animals in all groups successfully recognized the novel conspecific, as indicated by a marked increase in social interaction when the novel conspecific was introduced in trial 5, with no significant differences observed between treatment groups (Figure 4e-f). A social recognition index was calculated to compare the magnitude of recognition across groups; however, no significant between-group differences were observed (Figure 4h-j).

**Figure 4.**
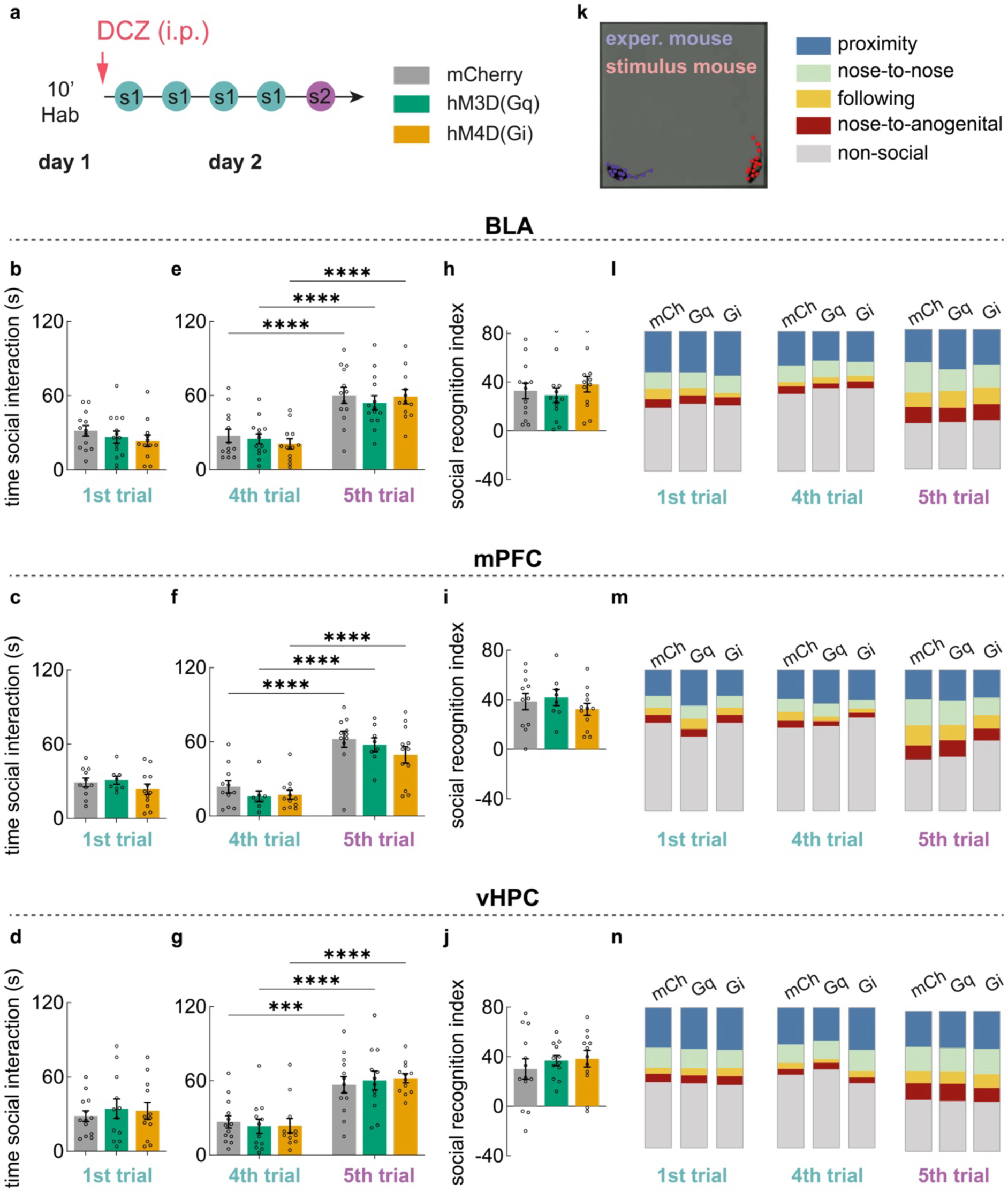
Sustained manipulation of VIP-expressing neurons does not significantly alter social behavioral measures. **a.** Schematic of the SRT, consisting of 1 day of chamber habituation followed by four 5-min presentations of Social Stimulus 1 and one 5-min presentation of Social Stimulus 2. **b.** Time spent in social interaction during the first trial following manipulation of VIP-expressing neurons in the BLA. **c.** Time spent in social interaction during the first trial following manipulation in the mPFC. **d.** Time spent in social interaction during the first trial following manipulation in the hippocampus. **e.** Time spent in social interaction during fourth and fifth trials in the BLA. **f.** Time spent in social interaction during fourth and fifth trials in the mPFC. **g.** Time spent in social interaction during fourth and fifth trials in the vHPC. **h.** social recognition index in the BLA, calculated as social interaction time during the fifth trial minus social interaction time during the fourth trial. **i.** social recognition index in the mPFC. **j.** social recognition index in the vHPC. **k.** Representative image of video-based automated mice recognition. **l.** Video-based automated classification of mouse behavior during first, fourth and fifth trials in the BLA. **m.** Video-based automated classification of mouse behavior during first, fourth and fifth trials in the mPFC. **n.** Video-based automated classification of mouse behavior during first, fourth and fifth trials in the vHPC. Data are expressed as the percentage of the first 2 min of each trial spent exhibiting each behavioral category. BLA, basolateral amygdala; mPFC, medial prefrontal cortex; SRT, social recognition task. Data are presented as mean ± SEM. Statistically significant post hoc comparisons are indicated by * p < 0.05, ** p < 0.01, *** p < 0.001, and **** p < 0.0001. Supplementary Table S5, S6 for statistical results.

To further characterize social interactions beyond total interaction time, an automated analysis pipeline was developed to classify and quantify the occurrence of specific behavioral elements. The pipeline successfully recognized and differentiate between the experimental mouse and social stimulus (example in Supplementary Figure S3). Therefore, several behavioral categories were defined: “proximity” and “nose-to-nose” behaviors, in which both animals contributed equally to the interaction; “following” and “nose-to-anogenital” behaviors, in which the experimental animal led the interaction; and “non-social” as the time spent not socially interacting (Figure 4k). While automated behavioral quantification using DeepLabCut revealed an expected increase in social behaviors such as “following” or “nose-to-anogenital” between trials 4 and 5 in all brain regions, as also observed in Figure 1l, sustained VIP modulation produced no significant differences between groups in the proportional breakdown of these specific behavioral categories (Figure 4 l-n, Supplementary Figure S5 l-m).

### Chemogenetic modulation of VIP^+^ neurons in mPFC impairs fear learning

Associative fear learning is highly conserved between rodents and humans, and dysregulated fear processing is a hallmark of psychiatric disorders such as post-traumatic stress disorder [40], making fear conditioning one of the most translational paradigms for dissecting the neural circuits of pathological fear. To investigate the role of VIP-expressing neurons in fear-related circuits, this population was chemogenetically modulated during an associative fear-conditioning paradigm (Figure 5a). In this paradigm, mice were exposed to a novel chamber (context A) on the training day, where they received five tone presentations (conditioned stimulus CS+) co-terminating with a 2 s, 0.4 mA foot shock (US), as well as five presentations of a different tone not paired with a foot shock (CS-). 24 h later, on the test day, mice were exposed to a distinct chamber (context B), where they received presentations of the same CS+ and CS- tones, but without the associated foot shock. During fear conditioning training on day 1, freezing increased across conditioning trials, with no significant differences between experimental groups following VIP-neuron modulation in the BLA, mPFC, or vHPC (Figure 5b-d). On the following day, memory retrieval was assessed during the test session. The first presentation of the CS+ in the absence of the aversive unconditioned stimulus (US) was used to probe fear-memory retrieval, whereas repeated presentations of the CS+ alone induce fear-memory extinction. No significant effects of chemogenetic modulation of BLA or vHPC VIP^+^ neurons were observed in baseline freezing (Figure 5e, g). In contrast, activation of mPFC VIP^+^ neurons significantly reduced baseline freezing relative to both the mCherry-control and hM4D(Gi) groups, although absolute freezing levels were close to zero in all groups (Figure 5f). During the first CS+ presentation on the test day, modulation of BLA or vHPC VIP^+^ neurons did not significantly affect freezing behavior (Figure 4h, j). In contrast, chemogenetic activation of mPFC VIP^+^ neurons pronouncedly reduced freezing relative to both the mCherry-control and hM4D(Gi) groups in the first CS+ presentation (Figure 4i). Together, these results indicate that sustained activation of VIP^+^ neurons in the mPFC during associative fear learning and memory retrieval selectively reduced fear expression, whereas comparable manipulation of VIP^+^ neurons in the BLA or vHPC did not produce detectable effects under the experimental conditions tested.

**Figure 5.**
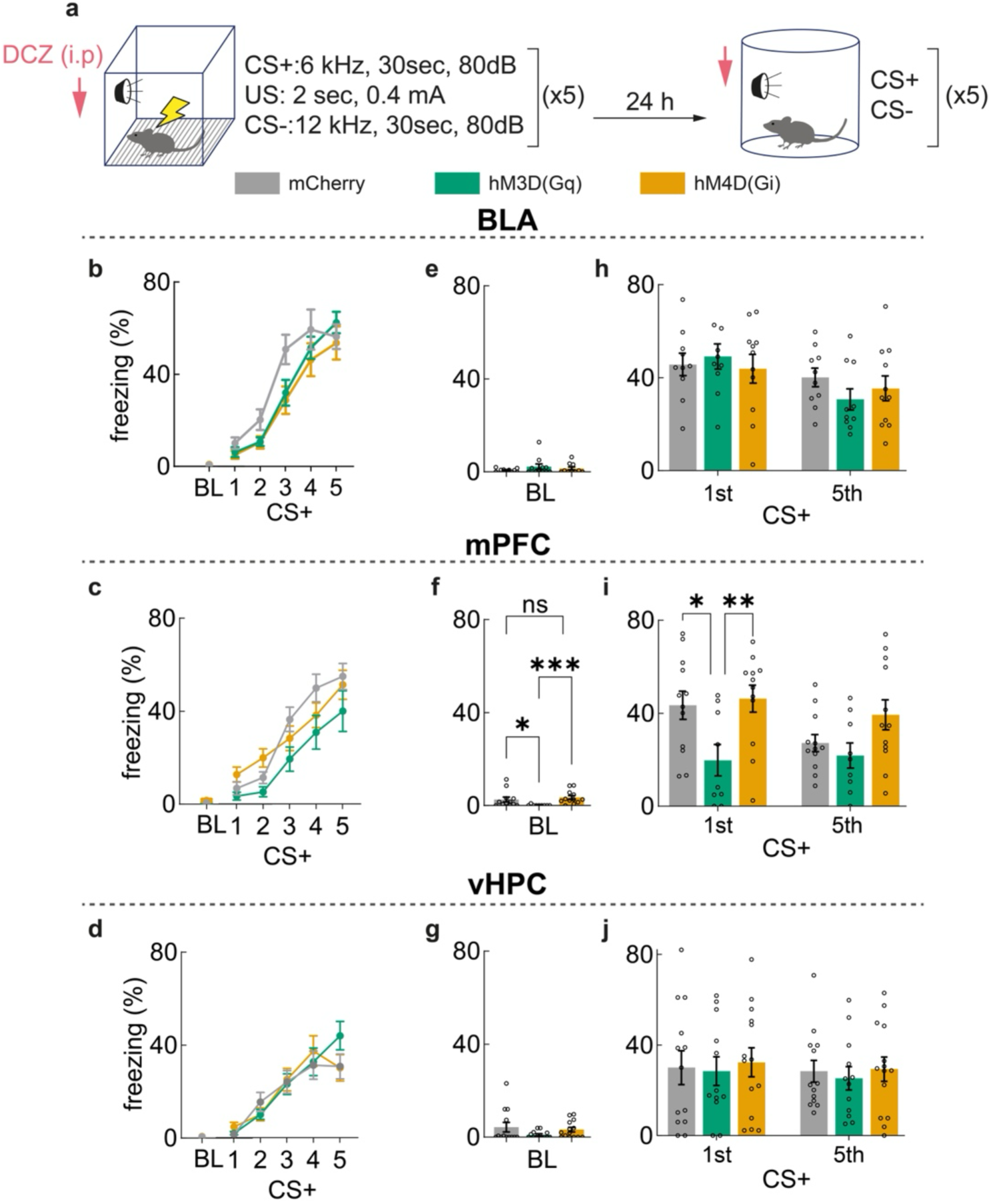
Sustained manipulation of mPFC VIP-expressing neurons significantly alters fear expression during memory retrieval. **a.** Schematic of the fear-conditioning and test paradigm. **b.** Freezing during baseline and successive CS+ presentations during fear conditioning following manipulation of VIP-expressing neurons in the BLA. **c.** Freezing during baseline and successive CS+ presentations during fear conditioning following manipulation in the mPFC. **d.** Freezing during baseline and successive CS+ presentations during fear conditioning following manipulation in the vHPC**. e.** Baseline freezing on test following manipulation in the BLA. **f.** Baseline freezing on test following manipulation in the mPFC. **g.** Baseline freezing on test following manipulation in the vHPC. **h.** Freezing during CS+ presentations on test following manipulation in the BLA. **i.** Freezing during CS+ presentations on test following manipulation in the mPFC. **j**. Freezing during CS+ presentations on test following manipulation in the vHPC. Data are presented as mean ± SEM. BLA, basolateral amygdala; mPFC, medial prefrontal cortex; vHPC, ventral hippocampus; CS+, conditioned stimulus paired with the unconditioned stimulus. Statistically significant post-hoc comparisons are indicated by * p < 0.05, ** p < 0.01, *** p < 0.001, and **** p < 0.0001. Supplementary Table S10, S11 for statistical results.

## Discussion

Understanding the role of distinct subtypes of GABAergic inhibitory neurons in the neuronal circuits that mediate psychiatrically relevant behaviors is essential for interpreting their contribution to the pathophysiology of psychiatric disorders. In the present study, we conducted a systematic analysis of the function of VIP^+^ GABAergic neurons in the mouse corticolimbic system in anxiety-related and social behaviors as well as in associative fear learning. Using an AAV-mediated chemogenetic approach, we show that sustained activation or inhibition of VIP^+^neurons in the BLA, mPFC or vHPC has no consequences for any of the anxiety-related or social behaviors assessed. In contrast, sustained activation of VIP^+^ neurons in the mPFC, but not the BLA or the vHPC, disrupted associative fear learning. Our study identifies regional and behavioral domain specificity of VIP^+^ neuron function and provides a framework for the study of VIP^+^ neuron dysfunction in mouse models of psychiatric and neurodevelopmental disorders. A central objective of our study was to provide a systematic and comparative assessment of the consequences of the disruption of VIP^+^ neuron function in the corticolimbic system on behavioral domains related to psychiatric disorders. Of particular interest in this respect are anxiety disorders, which are known to involve disruptions in GABAergic inhibitory transmission [3, 41]. One of the primary pharmacotherapies in use for the treatment of anxiety disorders are the benzodiazepines, which exert their anxiolytic effect through positive allosteric modulation of GABA_A_Rs [5, 6], indicating the importance of identifying the precise GABAergic mechanisms that contribute to anxiety behaviors. Indeed, an initial study reported that optogenetic inhibition of VIP^+^ neurons in the mouse mPFC reduced entries into the open arms of an EPM, but only if delivered specifically during arm entrance, not in response to sustained optogenetic inhibition throughout the behavioral paradigm [42]. In contrast, other anxiety-related behaviors remained unaffected in this study [42], consistent with a lack of effect of VIP^+^ neuron modulation in mPFC on anxiety behaviors in subsequent studies [43–45]. Similarly, chemogenetic or optogenetic inhibition of VIP^+^ neurons in vHPC did not affect anxiety-related behavioral outcomes [46, 47], although chemogenetic silencing of these neurons did attenuate the anxiogenic effect of a selective GABA_A_Rα5 modulator [47]. The results from the present study systematically extend these observations to demonstrate that anxiety-related behaviors are unaffected by either sustained activation or sustained inhibition of VIP^+^ neurons in either the mPFC, the BLA, or the vHPC. In combination with previous reports, our findings indicate that dysfunction of VIP^+^ GABAergic neurons in corticolimbic circuits is unlikely to be a driving force in the pathophysiology of anxiety disorders, and that therapeutic strategies targeting the GABAergic system may be more likely to act through other subcategories of inhibitory neurons.

A further behavioral domain which has previously received very little attention in the context of VIP^+^ neuron function are innate social behaviors. A limited number of studies have indicated potential roles of mPFC VIP^+^ neurons in individual aspects of social function. In particular, activation of VIP^+^ neurons in the mPFC was recently reported to induce winning behavior in a tube test of social dominance [44], while activation of VIP^+^ neurons selectively in the IL but not PreL reduced social interaction time between two mice [45]. Interestingly, while VIP^+^ neurons in the anterior cingulate cortex were shown to respond to social stimuli, the representation of these stimuli was highly unstable over time [48], raising questions on the role of these representations in social contexts. To our knowledge, no studies have reported any effects of VIP^+^ neurons in the mouse basolateral amygdala or ventral hippocampus in the regulation of social functions. Consistent with this notion, our systematic analysis revealed no effect of chemogenetic activation or inhibition of VIP^+^ neurons in any of the brain regions investigated on any of several parameters of social interest or social recognition analysed. Our findings lend important support to the conclusion that VIP^+^ neurons do not play a widespread role in the regulation of innate social behaviors throughout the corticolimbic system and therefore may not represent a key target for the study of social dysfunction in animal models of psychiatric and neurodevelopmental models.

In stark contrast to this very limited role for VIP^+^ neurons in the modulation of innate anxiety and social behaviors, a substantial and growing body of literature proposes a role for VIP^+^ neurons in regulating associative learning. Due to their morphological and functional properties in predominantly targeting other GABAergic inhibitory neurons, VIP^+^ neurons have been shown to contribute to disinhibitory circuit motifs in numerous contexts throughout the forebrain [12, 14, 49–52]. These disinhibitory circuit motifs have been demonstrated to play a key role in gating synaptic plasticity or stimulus discrimination during a wide range of associative learning paradigms [15, 17, 42, 46, 53–55], as well as motor learning [56, 57], spatial learning [18, 58, 59] and social memory [60]. Consistent with this notion of VIP^+^ neurons as an important plasticity gate, we report that sustained activation of VIP^+^ neurons in the mPFC disrupts behavioral performance in an auditory fear conditioning paradigm. The precise circuit mechanisms by which this disruption takes place remain to be explored. Given that chemogenetic activation of VIP^+^ neurons occurred during both acquisition and expression of the fear memory, we cannot at this point distinguish in which of the two phases of the paradigm VIP^+^ neurons exert their effect. Moreover, it will be interesting in the future to investigate how sustained chemogenetic activation of VIP^+^ neurons affects the disinhibitory motif proposed to the basis for the plasticity gate, and why sustained chemogenetic inhibition of the same neurons does not appear to detectably alter associative fear learning. Nevertheless, our findings clearly support the notion that disruption of VIP^+^ neuron function is most likely to influence behaviors that depend on plasticity mechanisms, rather than those that are innately hard-wired in neuronal circuits.

A second major objective of our study was to identify whether particular brain regions in the corticolimbic system may be more vulnerable to the sustained disruption of VIP^+^ neuron function than others. In this context, it is interesting to note that associative fear learning was affected by chemogenetic manipulation of VIP^+^ neurons only in the mPFC, but not in the BLA or vHPC, despite the fact that both are known to play key roles in mediating auditory fear conditioning [61–63]. The fact that the mPFC appears to be particularly vulnerable to the disruption of VIP^+^ neuron function may be unsurprising given that a wide range of resulting behavioral phenotypes have been previously reported, including alterations in auditory discrimination [14], goal-directed behavior [64], inhibitory control [43, 65], attentional function [16], cognitive flexibility [66], regulation of food intake [45], pain sensitivity [67], and spatial working memory [59]. Perhaps more surprising is the lack of a phenotype of VIP^+^ neurons in the BLA and vHPC, particularly given that in both brain regions, optogenetic experiments have shown that temporary, stimulus-locked manipulations of VIP^+^ neurons do modulate fear learning [46, 54]. It should be noted that extremely limited information is available on the consequences of manipulating VIP^+^ neurons in the BLA on any behavioral outcomes, with only a single currently available study, which demonstrates that optogenetic inhibition of VIP^+^ neurons in the BLA selectively during presentation of the US impairs auditory fear learning [54]. It is conceivable that, as was reported for the role of mPFC VIP^+^ neurons in anxiety behaviors [42], temporary stimulus-specific manipulations may have distinct consequences from sustained manipulations across the duration of the behavioral paradigm, as would be the case for our chemogenetic activation or inhibition. The same logic may apply to the vHPC, where a previous study demonstrated that optogenetic inhibition of VIP^+^ neurons during presentation of the CS, trace and US of a trace fear conditioning paradigm impairs formation of a CS-US association [46]. In the latter case, an additional difference may result from the behavioral paradigm used, since trace fear conditioning may involve distinct circuits compared to the delay fear conditioning paradigm applied in our study, although both are known to depend on the vHPC [61, 63]. Again, very few other studies are available that investigate the consequences of selective manipulation of VIP^+^ neuron activity in vHPC on any behavioral output, highlighting the urgent need for additional studies investigating the detailed role of these neurons in both vHPC and BLA. Nevertheless, our findings, in combination with the available literature, would indicate that sustained disruptions of VIP^+^ activity are more likely to affect behaviors dependent on the mPFC than the BLA or vHPC.

The ultimate rationale for our study was to provide a basis for the future study of VIP^+^ neuron dysfunction in animal models of psychiatric and neurodevelopmental disorders. Alterations in the function of VIP^+^ neurons have been observed in a number of mouse models, including of general neurodevelopmental delay [19], Dravet syndrome [20, 21], Rett syndrome [22], Fragile X syndrome [23], 22q11 deletion syndrome [24], and epilepsy [25]. Together with other published findings, our results imply that alterations in VIP^+^ neuron function resulting from genetic mutations may be particularly likely to disrupt mPFC-mediated plasticity-dependent learning processes. In addition, the discrepancies observed between reported results from temporary, stimulus-locked optogenetic manipulations and our findings using sustained chemogenetic manipulations highlight the challenges in transferring knowledge from studies primarily designed to acutely elucidate precise encoding mechanisms and circuit motifs to a disease-related setting with persistent functional disruption. These observations are particularly relevant in light of the recent notion that AAV-mediated chemogenetic approaches may be used as direct therapeutic strategies in neurodevelopmental and psychiatric disorders [27–29]. Ultimately, selective disruption of the genes of interest in VIP^+^ neurons and detailed analysis of the consequences on VIP^+^ synapses and circuits in the mPFC will be necessary to fully understand the mechanisms by which pathophysiological mutations may result in psychiatrically relevant cognitive impairments.

## Acknowledgments

This work was supported by the Deutsche Forschungsgemeinschaft (Heisenberg grant KR5239/1 and grant KR5239/2, both to D.K.-B.), as well as by a Ph.D. fellowship from the Focus Program Translational Neurosciences at the Johannes Gutenberg University Mainz (to M.T.). J.H. and M.A.N. were funded by the Boehringer Ingelheim Foundation. M.J.S. was supported by the Werner Reichenberger Foundation. The authors are grateful to Mikkel Blake, Alexander Wenzel, Christine Oswald and the staff at the Translational Animal Research Center for excellent technical support and animal care, and to Sebastian Wiens for video processing.

## Author contributions

D.K.-B. and M.T. conceived the study and designed all experiments, with the assistance of D.P.C. M.T. performed all surgical, pharmacological and behavioral procedures and analysed behavioral data, under the supervision of D.K.-B. and D.P.C. M.T. performed all immunohistochemical analysis, with the assistance of Y.G. M.A.N. developed algorithms for the automated analysis of social behaviors using DeepLabCut under the supervision of J.H., and results derived from the automated social behavior analysis will be included in her MD thesis. D.K.-B., J.H. and M.J.S. provided supervision, funding and equipment. M.T. and D.K.-B. prepared the manuscript, and all authors edited the manuscript and approved the final version.

