## Supplementary Information for "Consequences of sustained modulation of VIP-expressing neurons in mouse corticolimbic regions on anxiety-related avoidance behaviors, social functioning, and associative fear learning"

### SUPPLEMENTARY METHODS FOR AUTOMATED BEHAVIORAL PROFILING

Objective quantification of mouse behavior was performed using video-based automated analysis. Body part tracking was carried out with DeepLabCut [1]; pose estimation data was stored per video in HDF5 format, containing the pixel coordinates of tracked body parts for the two mice across all frames. Following the DeepLabCut multi-animal guidelines [2], body parts with a DeepLabCut-assigned likelihood of $p < 0.6$ were classified as missing. Custom Python-based scripts were applied for improved tracking and behavioral classification. All post-processing of the DeepLabCut output and behavioral analyses were implemented as custom Python (version 3.8.10) scripts using NumPy [3], Pandas [4], SciPy [5], and OpenCV [6].

We trained a multi-animal DeepLabCut model based on the ResNet-50 architecture on our video recordings [2]. The annotated body landmarks were selected according to the body-part scheme of DeepOF [7]. While the DeepLabCut multi-animal framework provides reliable pose estimation, close physical interactions and occlusions can interfere with body-part association and identity tracking [2]. In our recordings, these situations require additional post-processing.

#### 1. Correction of mouse identities

Identity switches occur when animals are in close proximity and involve either a "jump" of a body part from one animal to the other, or the disappearance of one animal from the tracking, see Supplementary Figure S3. These criteria identified potential identity switches, after which we reassigned identity.

***Potential identity switches*** were identified as a low inter-animal distance < 60 pixels (7.72 cm), calculated as the Euclidean distance between the *Center* body parts of the two mice, and one of the following conditions:

1) Loss of tracking: The first frame following the loss of all tracked body parts of one mouse identified the time point of a potential identity switch.
2) A "jump" in body-part location: The body-part displacement was calculated as the Euclidean distance of the body part location of two subsequent frames. The mean body-part displacement over all tracked body parts summed for both mice was defined as the overall body-part displacement. The frame following the largest overall body-part displacement within the period of close proximity identified the time point of a potential identity switch.

***Assignment of animal identity*** was based on the lighter color of the shaved back of the social stimulus mouse and its smaller size due to its younger age. As the social stimulus mouse was shaved on its back, dorsal brightness was compared between both mice as distinguishing feature. The dorsal brightness for both animals was extracted from the value channel (brightness) of the HSV color space around the *Spine_2* body part, as mean over a radius of 3 pixels (~4 mm). A lower dorsal brightness identified the experimental mouse with dark dorsal fur. When differences in dorsal brightness were not clearly detectable, body size was compared between the mice as supplementary feature. Body size was calculated as the area of a polygon defined by the following eight body parts: *Nose, Right_ear, Right_fhip, Right_bhip, Tail_base, Left_bhip, Left_fhip, and Left_ear*. A larger body size identified the older experimental mouse.

**2. Post-processing of DeepLabCut Pose Estimations**

Missing body-part coordinates were linearly interpolated when the distance between the last and the next valid detection was less than 60 pixels (7.72 cm), corresponding approximately to the body length of a mouse. For the key body landmarks (*Nose*, *Center* and *Tail_base*) of both animals, missing detections spanning distances greater than 60 pixels were manually corrected using a custom Python-based annotation interface. To reduce frame-to-frame tracking noise, all body-part trajectories were smoothed using a one-dimensional Gaussian filter (scipy.ndimage.gaussian_filter1d, σ = 2). The resulting post-processed DeepLabCut pose estimation was used to classify behavioral events.

**3. Python-Based Classification of Social Behaviors**

The pose estimation data was categorized into five behaviours, time points were assigned to the categories in the following order:

1. A *following* region of interest (ROI) was defined as the combined area of 25-px-radius (3.22 cm) circles centred on the stimulus mouse's *Tail_base* over the preceding 10 frames (≈0.67 s). The behaviour *following* was identified when the *Nose* of the experimental mouse entered this ROI while both animals were oriented in a similar direction, i.e. body orientation differed by less than 45°, and while the mouse was moving, i.e. the *Nose* was moving at a speed of 75 pixels/s and its speed remains above 39 pixels/s for the following 10 frames (≈0.67 s).
2. The behaviour *nose-to-anogenital* was identified when the distance between the *Nose* of the experimental animal and the *Tail_base* of the stimulus mouse was smaller than 20 pixels (< 2.57 cm).
3. The behaviour *nose-to-nose* was identified when the distance between the noses (body part *Nose*) of both animals was smaller than 20 pixels (< 2.57 cm).
4. The remaining social behavior was summarized as *proximity*, identified by a distance between the *Center* body parts of both mice smaller than 60 pixels (< 7.72 cm), or when the stimulus mouse was following the experimental mouse, i.e. the *Nose* of the stimulus animal and *Tail_base* of the experimental mouse were closer than 25 pixels (< 3.22 cm) for at least 10 frames (≈ 0.67 s); and body orientation differed by less than 45°. When the *Nose* of one mouse is simultaneously reported as close to the *Nose* and *Tail_base* of the other mouse, or for small *Nose*-to-*Tail_base* distances for both mice, the time points are recategorized as *proximity*.
5. Remaining time points were categorized as *non-social*.

Due to noise in the tracking, following of the experimental mouse by the stimulus mouse and the inverse, following of the stimulus mouse by the experimental mouse, was detected simultaneously for short durations, typically less than 0.3 seconds. The time points were recategorized as *nose-to-anogenital* if the corresponding condition was matched, as *proximity* if the *Center* distance was less than 60 pixels, and otherwise as *non-social*.

### SUPPLEMENTARY FIGURES

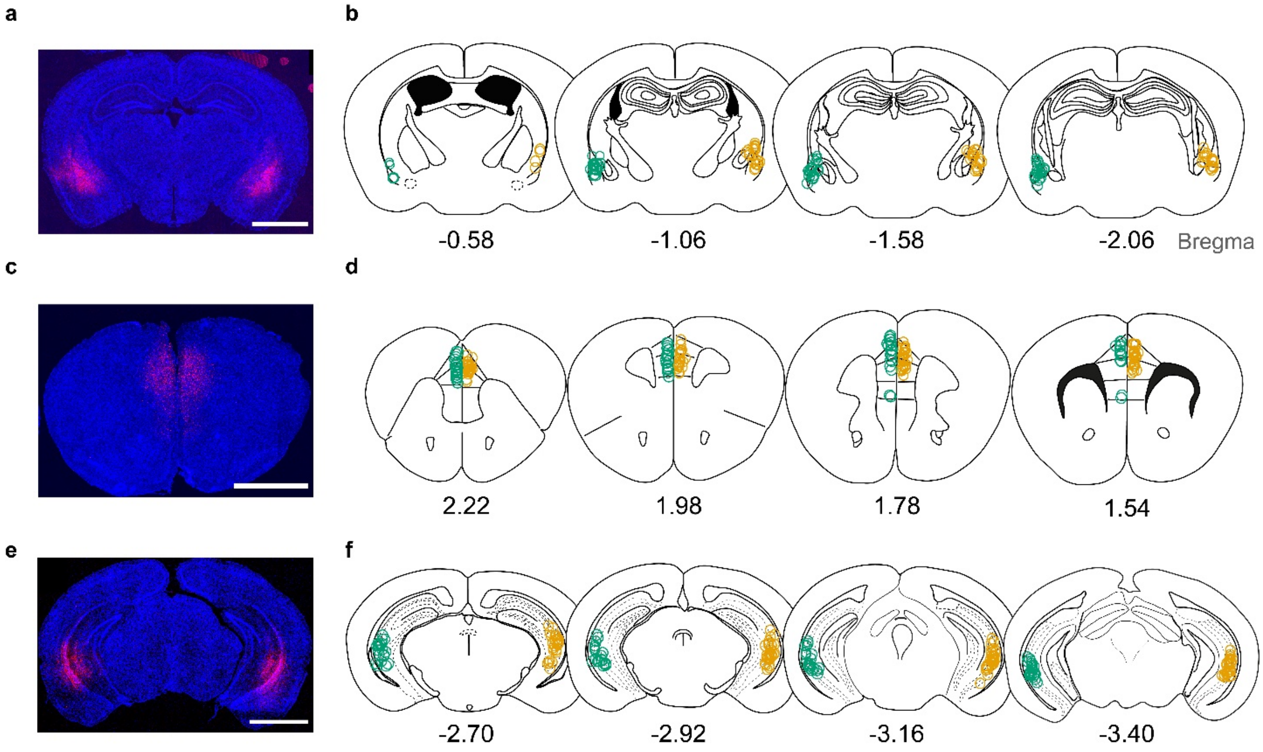


**Supplementary Figure S1. Anatomical validation of DREADD expression in targeted corticolimbic regions. (a)** Representative confocal image of DREADD expression in the basolateral amygdala (BLA). **(b)** Coronal maps showing the center of DREADD expression in the BLA across all animals included in statistical analyses. **(c)** Representative confocal image of DREADD expression in the medial prefrontal cortex (mPFC). **(d)** Coronal maps showing the center of DREADD expression in the mPFC across all animals included in statistical analyses. **(e)** Representative confocal image of DREADD expression in the ventral hippocampus (vHPC). **(f)** Coronal maps showing the center of DREADD expression in the vHPC across all animals included in statistical analyses. hM3D(Gq), green; hM4D(Gi), orange. Scale bar, 2 mm.

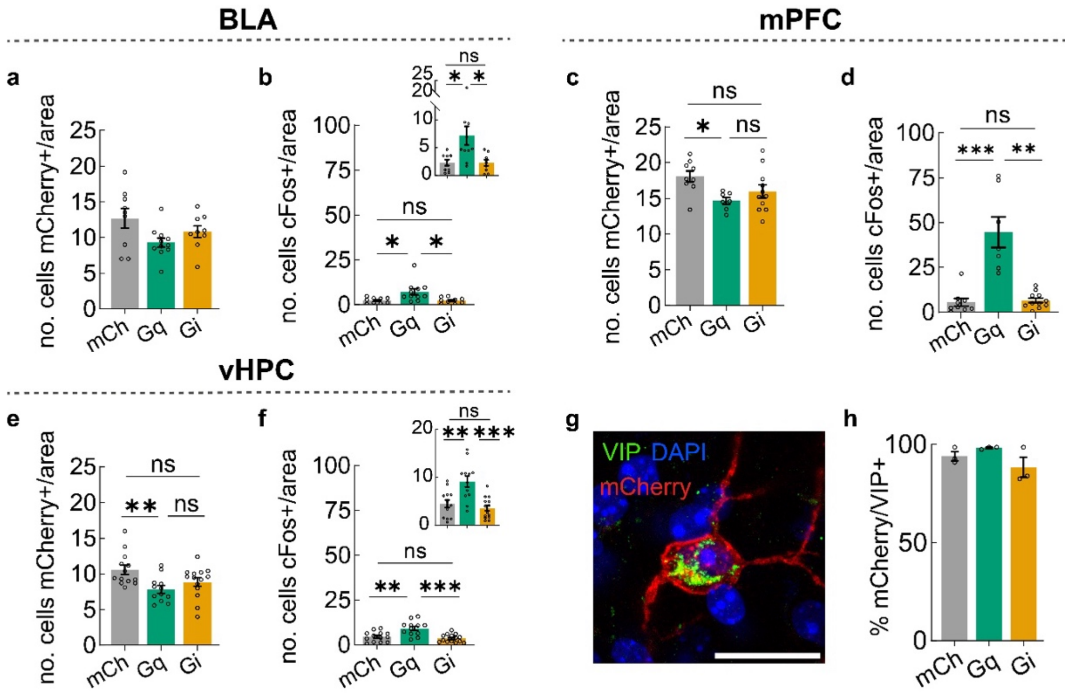


**Supplementary Figure S2. Validation of DCZ-induced activation of DREADD-expressing VIP neurons across corticolimbic regions. (a)** Number of mCherry+ cells per 0.34 mm² in the basolateral amygdala (BLA). **(b)** Number of cFos+ cells per 0.20 mm² in the BLA. **(c)** Number of mCherry+ cells per 0.20 mm² in the medial prefrontal cortex (mPFC). **(d)** Number of cFos+ cells per 0.34 mm² in the mPFC. **(e)** Number of mCherry+ cells per 0.34 mm² in the ventral hippocampus (vHPC). **(f)** Number of cFos+ cells per 0.34 mm² in the vHPC **(g)** Representative confocal image showing VIP immunostaining colocalized within an mCherry-expressing cell **(h)** Percentage of mCherry+ cells showing VIP immunoreactivity. Scale bar 20 µm. Data are presented as mean ± SEM. Statistically significant post-hoc analyses are marked with *p<0.05, ** p<0.01, *** p<0.001, *****p<0.0001. Supplementary Table S3 for statistical results.

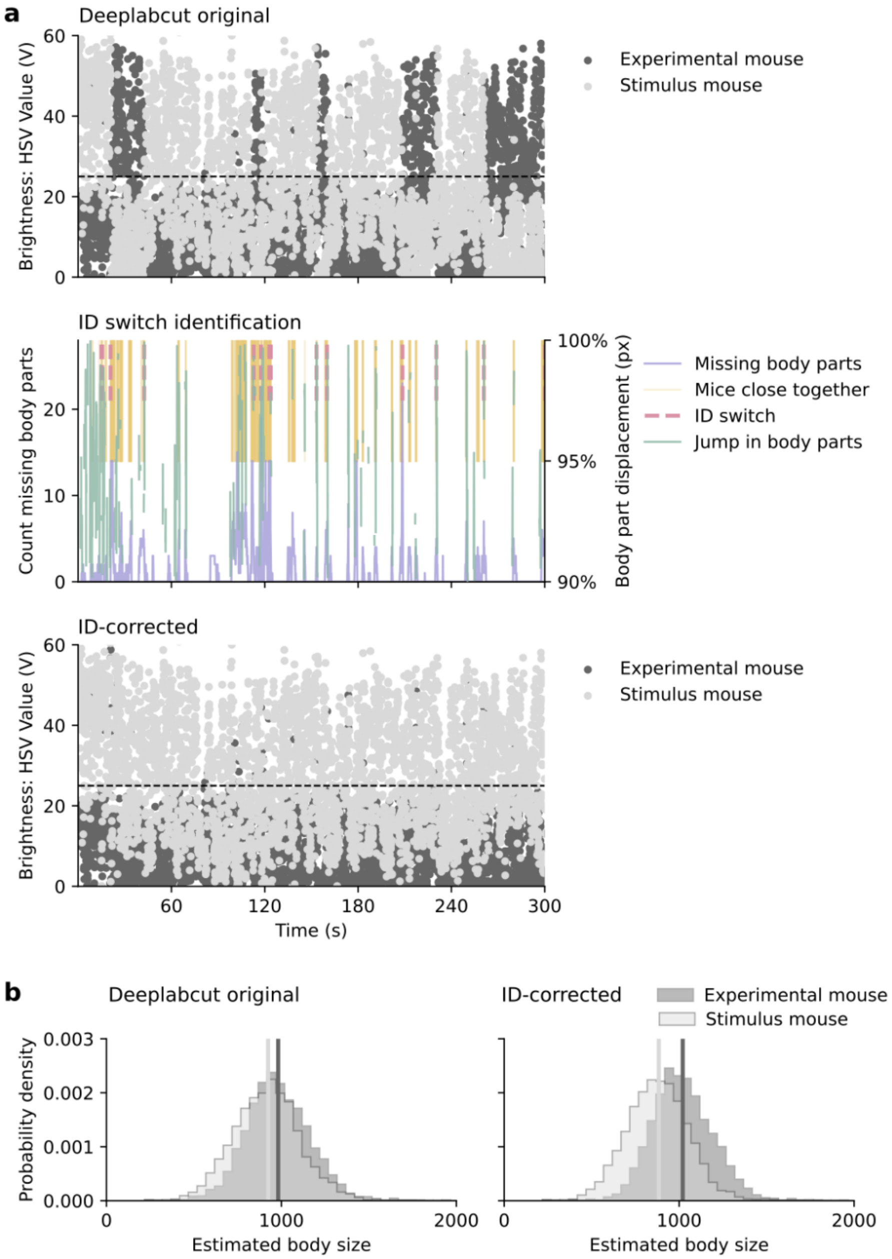


**Supplementary Figure S3. Identity correction exemplified for one mouse and one SRT session. (a)** Top: Before identity correction, higher dorsal brightness, which marks the shaved stimulus mouse, switches between the two mice. Middle: These switches occur only when animals are in close proximity (pink shading) and either the number of tracked body parts reduces because one animal disappears from the tracking (blue line), or when body parts jump between frames (high velocity body parts in green, right axis). Bottom: After identity correction, dorsal brightness is mostly consistent with animal identity. (**b)** Before identity correction, the body size of both animals is similar (left, 6% difference in mean size). After identity correction, the experimental mouse has a higher body size, as expected due to its older age (right, 15% difference in mean size).

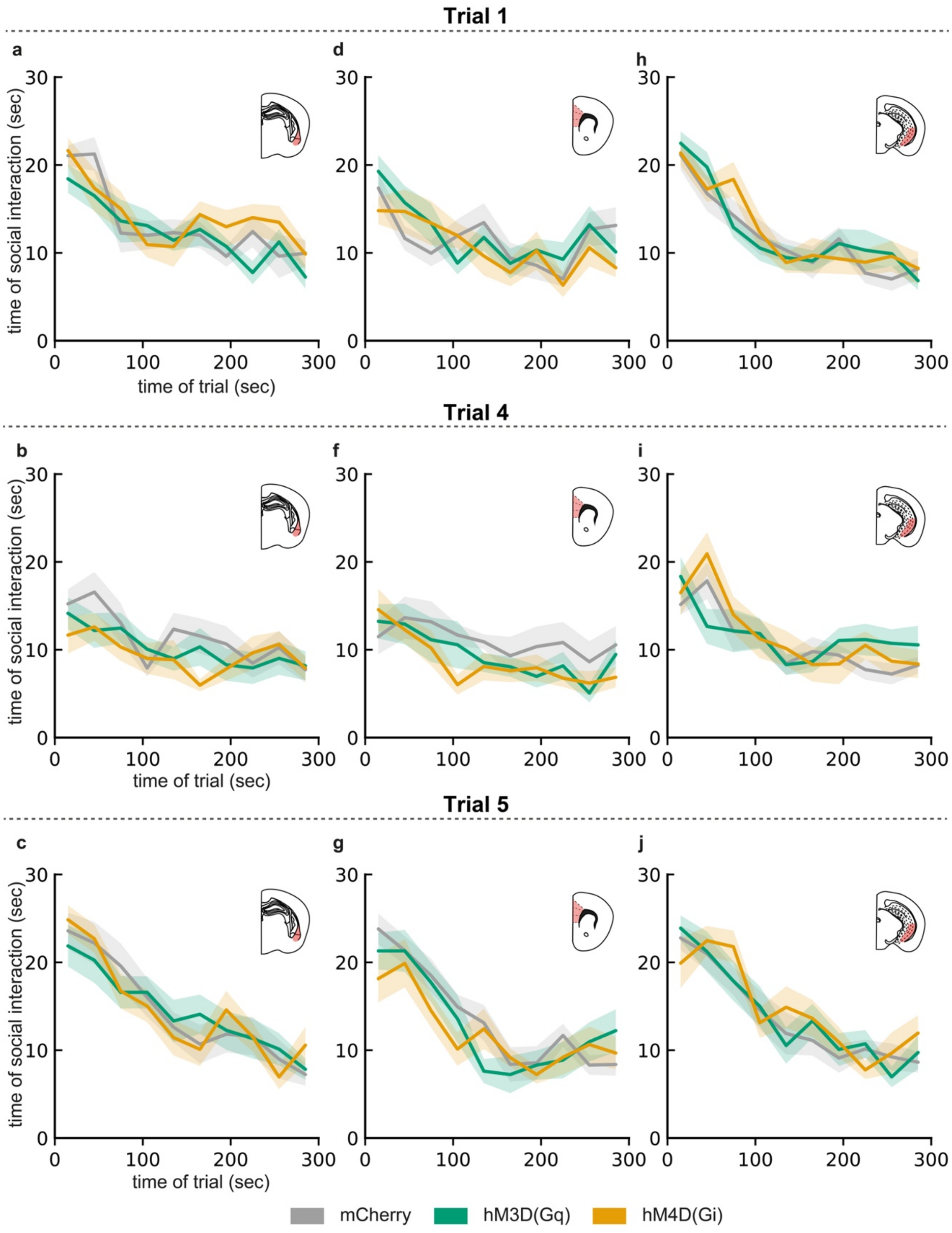


**Supplementary Figure S4. Time course of social interaction during the first 2 min of the Social Recognition Task. (a–c) BLA**: time course of social interaction during trial 1 **(a),** trial 4 **(b),** and trial 5 **(c)** in animals with DREADD expression in the basolateral amygdala (BLA). **(d–f) mPFC**: time course of social interaction during trial 1 **(d)**, trial 4 **(e)**, and trial 5 **(f)** in animals with DREADD expression in the medial prefrontal cortex (mPFC). **(g–i)** vHPC: time course of social interaction during trial 1 **(g)**, trial 4 **(h)**, and trial 5 **(i)** in animals with DREADD expression in the ventral hippocampus (vHPC).

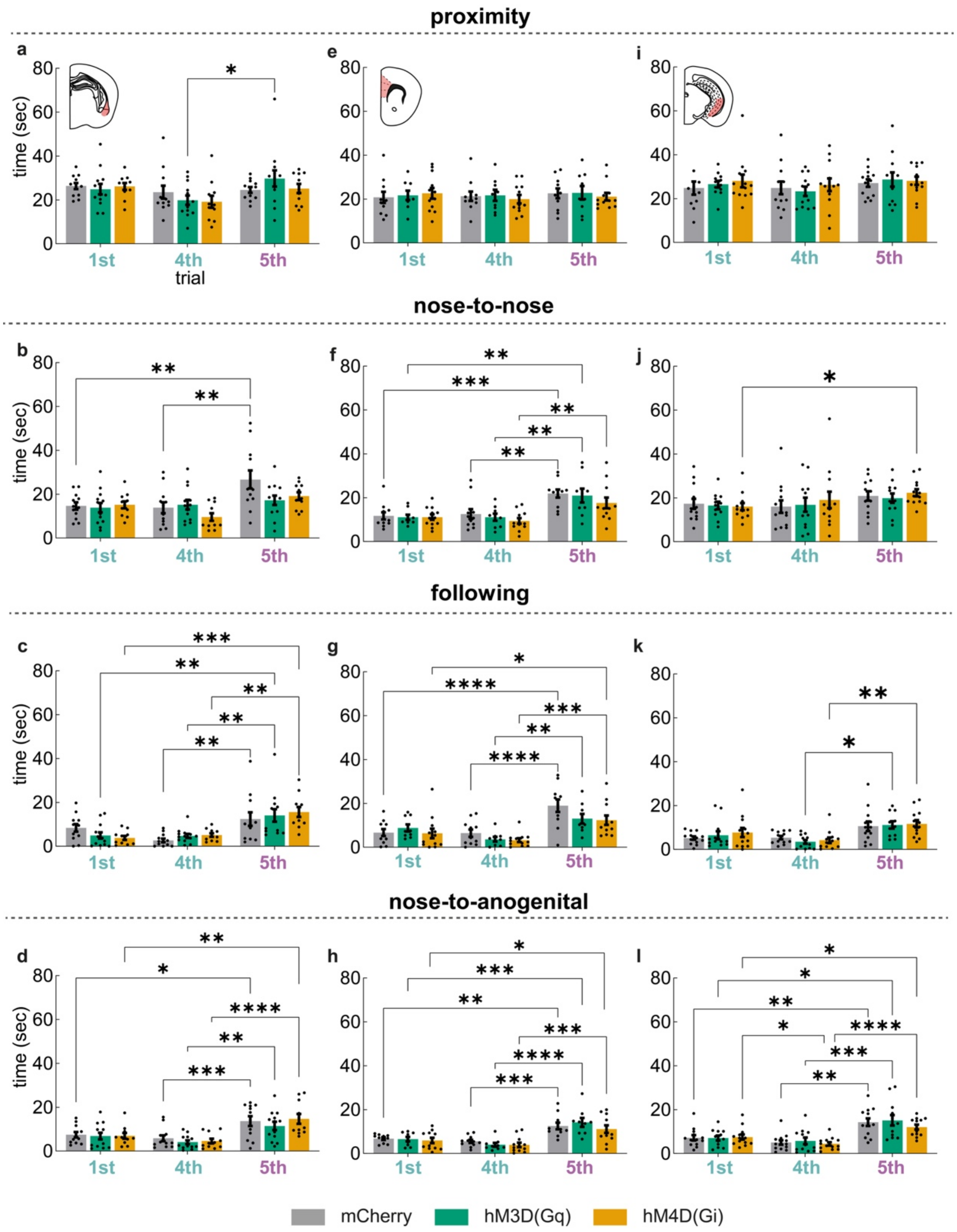


**Supplementary Figure S5. Detailed occurrence of specific behavioral elements during the Social Recognition Task (SRT). (a–d) BLA**: time spent performing "proximity" (a), "nose-to-nose poke" (b), "following" (c), and "nose-to-anogenital poke" (d) during the first 2 min of the SRT in animals with DREADD expression in the basolateral amygdala (BLA). **(e–h) mPFC**: time spent performing "proximity" (e), "nose-to-nose poke" (f), "following" (g), and "nose-to-anogenital poke" (h) during the first 2 min of the SRT in animals with DREADD expression in the medial prefrontal cortex (mPFC). **(i–l) vHPC**: time spent performing "proximity" (i), "nose-to-nose poke" (j), "following" (k), and "nose-to-anogenital poke" (l) during the first 2 min of the SRT in animals with DREADD expression in the vHPC. Data are presented as mean ± SEM. Statistically significant post-hoc analyses are marked with *p<0.05, ** p<0.01, *** p<0.001, *****p<0.0001. Supplementary Table S7, S8, S9 for statistical results.

### SUPPLEMENTARY TABLES

**Supplementary Table S1. Statistical comparisons of DCZ off-target effects on behavior.** Group differences were assessed using an unpaired t-test (parametric data) or Mann-Whitney U test (non-parametric data), as appropriate.

| | Distribution | p-value | Significance |
|---|---|---|---|
| Entries open arms | Normal | 0.245 | ns |
| Time open arms | Not normal | 0.721 | ns |
| Entries center zone | Not normal | 0.778 | ns |
| Time in center zone | Normal | 0.633 | ns |
| Total distance travelled | Normal | 0.706 | ns |
| Social interaction time at 1st trial | Normal | 0.279 | ns |
| Social Recognition Index | Normal | 0.341 | ns |
| Baseline fear conditioning | Not normal | 0.902 | ns |
| Baseline text/extinction | Not normal | 0.771 | ns |

**Supplementary Table S2. Statistical comparisons of DCZ off-target effects on behavior.** Group differences were assessed using RM two-way ANOVA.

| | Trial x Group | | Group | | Trial | | Post-hoc |
|---|---|---|---|---|---|---|---|
| | F-value | p-value | F-value | p-value | F-value | p-value | Adjusted p-value |
| Time spent in interaction (t4, t5) | $F_{(1, 13)}$ = <1 | 0.341 | $F_{(1, 13)}$ = <1 | 0.982 | $F_{(1, 13)}$ = 43.55 | <0.0001 | Saline: 0.0003<br>DCZ: 0.0025 |
| Fear conditioning | $F_{(4, 56)}$ = 1.157 | 0.340 | $F_{(1, 14)}$ = <1 | 0.602 | $F_{(2.673, 37.42)}$ = 25.32 | <0.0001 | Continue on table below |
| Test/extinction | $F_{(4, 56)}$ = 1.110 | 0.361 | $F_{(1, 14)}$ = <1 | 0.799 | $F_{(2.801, 39.21)}$ = 1.511 | 0.2283 | - |

**Supplementary Table S3. Statistical comparisons of Validation of DCZ-induced activation of DREADD-expressing VIP neurons across corticolimbic regions.** Group differences were assessed using a one-way ANOVA (parametric data) or Kruskal-Wallis test (non-parametric data), as appropriate.

| | Distribution | p-value | Significance | Post-hoc |
|---|---|---|---|---|
| **BLA:** % mCherry$^+$/cFos$^+$ | Not normal | <0.0001 | **** | mCh vs. Gq = 0.002<br>mCh vs. Gi = 0.945<br>Gq vs. Gi = <0.0001 |
| **mPFC:** % mCherry$^+$/cFos$^+$ | Not normal | 0.0002 | *** | mCh vs. Gq = 0.018<br>mCh vs. Gi = 0.505<br>Gq vs. Gi = 0.0001 |
| **vHPC:** % mCherry$^+$/cFos$^+$ | Not normal | <0.0001 | **** | mCh vs. Gq = 0.010<br>mCh vs. Gi = 0.106<br>Gq vs. Gi = <0.0001 |

| | Distribution | p-value | Significance | Post-hoc |
|---|---|---|---|---|
| **BLA:** mCherry$^+$/area | Normal | 0.057 | ns | - |
| **BLA:** cFos$^+$ /area | Not normal | 0.005 | ** | mCh vs. Gq = 0.019<br>mCh vs. Gi = >0.9999<br>Gq vs. Gi = 0.012 |
| **mPFC:** mCherry$^+$/area | Normal | 0.03 | * | mCh vs. Gq = 0.027<br>mCh vs. Gi = 0.155<br>Gq vs. Gi = 0.51 |
| **mPFC:** cFos$^+$ /area | Not normal | 0.0004 | *** | mCh vs. Gq = 0.001<br>mCh vs. Gi = >0.9999<br>Gq vs. Gi = 0.004 |
| **vHPC:** mCherry$^+$/area | Normal | 0.009 | ** | mCh vs. Gq = 0.007<br>mCh vs. Gi = 0.100<br>Gq vs. Gi = 0.439 |
| **vHPC:** cFos$^+$ /area | Normal | <0.0001 | **** | mCh vs. Gq = 0.001<br>mCh vs. Gi = 0.729<br>Gq vs. Gi = 0.0001 |

**Supplementary Table S4. Statistical comparisons of the effects of sustained modulation of VIP$^+$ neurons across corticolimbic regions on anxiety-like behavior.** Group differences were assessed using a one-way ANOVA (parametric data) or Kruskal-Wallis test (non-parametric data), as appropriate.

| | Distribution | p-value | Significance |
|---|---|---|---|
| **BLA:** Time in open arms | Normal | 0.944 | ns |
| **BLA:** Time in center zone | Not normal | 0.431 | ns |
| **BLA:** Total distance travelled | Normal | 0.95 | ns |
| **mPFC:** Time in open arms | Normal | 0.153 | ns |
| **mPFC:** Time in center zone | Not normal | 0.975 | ns |
| **mPFC:** Total distance travelled | Normal | 0.939 | ns |
| **vHPC:** Time in open arms | Normal | 0.581 | ns |
| **vHPC:** Time in center zone | Normal | 0.689 | ns |
| **vHPC:** Total distance travelled | Normal | 0.488 | ns |

**Supplementary Table S5. Statistical comparisons of the effects of sustained modulation of VIP+ neurons across corticolimbic regions on social behavior.** Group differences were assessed using a one-way ANOVA (parametric data).

| | Distribution | p-value | Significance |
|---|---|---|---|
| **BLA:** Time of social interaction in 1st trial | Normal | 0.480 | ns |
| **BLA:** Social recognition index | Normal | 0.609 | ns |
| **mPFC:** Time of social interaction in 1st trial | Normal | 0.391 | ns |
| **mPFC:** Social recognition index | Normal | 0.533 | ns |
| **vHPC:** Time of social interaction in 1st trial | Normal | 0.798 | ns |
| **vHPC:** Social recognition index | Normal | 0.644 | ns |

**Supplementary Table S6. Statistical comparisons of the effects of sustained modulation of VIP+ neurons across corticolimbic regions on social behavior.** Group differences were assessed using a RM two-way ANOVA.

| | Trial x Group | | Group | | Trial | | Post-hoc |
|---|---|---|---|---|---|---|---|
| | F-value | p-value | F-value | p-value | F-value | p-value | Adjusted p-value |
| **BLA:** Time of social interaction (t4, t5) | $F_{(2, 36)}$ = < 1 | 0.602 | $F_{(2, 36)}$ = < 1 | 0.742 | $F_{(1, 36)}$ = 85.38 | <0.0001 | mCh vs. Gq = <0.0001<br>mCh vs. Gi = <0.0001<br>Gq vs. Gi =<0.0001 |
| **mPFC:** Time of social interaction (t4, t5) | $F_{(2, 28)}$ = < 1 | 0.533 | $F_{(2, 28)}$ = 1.190 | 0.319 | $F_{(1, 28)}$ = 116.5 | <0.0001 | mCh vs. Gq = <0.0001<br>mCh vs. Gi = <0.0001<br>Gq vs. Gi =<0.0001 |
| **vHPC:** Time of social interaction (t4, t5) | $F_{(2, 34)}$ = < 1 | 0.642 | $F_{(2, 34)}$ = < 1 | 0.984 | $F_{(1, 34)}$ = 82.44 | <0.0001 | mCh vs. Gq = 0.0002<br>mCh vs. Gi = <0.0001<br>Gq vs. Gi =<0.0001 |

**Supplementary Table S7. Statistical comparisons of the effects of sustained modulation of BLA VIP+ neurons on different elements of social behavior.** Group differences were assessed using a RM two-way ANOVA.

| | Trial x Group | | Group | | Trial | | Post-hoc |
|---|---|---|---|---|---|---|---|
| | F-value | p-value | F-value | p-value | F-value | p-value | Adjusted p-value |
| **BLA:** Time spent "proximity" | $F_{(4, 66)}$ = 1.179 | 0.328 | $F_{(2, 33)}$ = < 1 | 0.78 | $F_{(2, 66)}$ = 5.180 | 0.008 | mCh<br>1st vs 4th =>0.999<br>1st vs 5th =>0.999<br>4th vs 5th =>0.999<br>Gq<br>1st vs 4th =>0.999<br>1st vs 5th =>0.999<br>4th vs 5th =0.024<br>Gi<br>1st vs 4th =0.425<br>1st vs 5th =>0.999<br>4th vs 5th =0.756 |
| **BLA:** Time spent "nose-to-nose" | $F_{(4, 66)}$ = 2.209 | 0.078 | $F_{(2, 33)}$ = 1.893 | 0.167 | $F_{(2, 66)}$ = 10.65 | <0.0001 | mCh<br>1st vs 4th =>0.999<br>1st vs 5th =0.003<br>4th vs 5th =0.001<br>Gq<br>1st vs 4th =>0.999<br>1st vs 5th =>0.999<br>4th vs 5th =>0.999<br>Gi<br>1st vs 4th =0.921<br>1st vs 5th =>0.999<br>4th vs 5th =0.055 |
| **BLA:** Time spent "following" | $F_{(4, 66)}$ = 1.531 | 0.203 | $F_{(2, 33)}$ = < 1 | 0.98 | $F_{(2, 66)}$ = 26.92 | <0.0001 | mCh<br>1st vs 4th =0.245<br>1st vs 5th =0.993<br>4th vs 5th =0.002<br>Gq<br>1st vs 4th =>0.999<br>1st vs 5th =0.003<br>4th vs 5th =0.002<br>Gi<br>1st vs 4th =>0.999<br>1st vs 5th =0.0002<br>4th vs 5th =0.001 |
| **BLA:** Time spent "nose-to-anogenital" | $F_{(4, 66)}$ = 0.457 | 0.767 | $F_{(2, 33)}$ = < 1 | 0.604 | $F_{(2, 66)}$ = 31.75 | <0.0001 | mCh<br>1st vs 4th =>0.999<br>1st vs 5th =0.016<br>4th vs 5th =0.001<br>Gq<br>1st vs 4th =>0.999<br>1st vs 5th =0.127<br>4th vs 5th =0.001<br>Gi<br>1st vs 4th =>0.999<br>1st vs 5th =0.002<br>4th vs 5th =<0.0001 |

**Supplementary Table S8. Statistical comparisons of the effects of sustained modulation of mPFC VIP+ neurons on different elements of social behavior.** Group differences were assessed using a RM two-way ANOVA.

| | Trial x Group | | Group | | Trial | | Post-hoc |
|---|---|---|---|---|---|---|---|
| | F-value | p-value | F-value | p-value | F-value | p-value | Adjusted p-value |
| **mPFC:** Time spent "proximity" | $F_{(4, 58)}$ = < 1 | 0.889 | $F_{(2, 29)}$ = < 1 | 0.937 | $F_{(2, 58)}$ = < 1 | 0.759 | - |
| **mPFC:** Time spent "nose-to-nose" | $F_{(4, 58)}$ = < 1 | 0.807 | $F_{(2, 29)}$ = 1.07 | 0.358 | $F_{(2, 58)}$ = 27.51 | <0.0001 | mCh<br>1st vs 4th =>0.999<br>1st vs 5th =0.001<br>4th vs 5th =0.002<br>Gq<br>1st vs 4th =>0.999<br>1st vs 5th =0.003<br>4th vs 5th =0.004<br>Gi<br>1st vs 4th =>0.999<br>1st vs 5th =0.053<br>4th vs 5th =0.006 |
| **mPFC:** Time spent "following" | $F_{(4, 58)}$ = 1.91 | 0.122 | $F_{(2, 29)}$ = 1.435 | 0.254 | $F_{(2, 58)}$ = 36.22 | <0.0001 | mCh<br>1st vs 4th =>0.999<br>1st vs 5th =0.0001<br>4th vs 5th =0.0001<br>Gq<br>1st vs 4th =0.281<br>1st vs 5th =0.710<br>4th vs 5th =0.002<br>Gi<br>1st vs 4th =>0.999<br>1st vs 5th =0.04<br>4th vs 5th =0.0003 |
| **mPFC:** Time spent "nose-to-anogenital" | $F_{(4, 58)}$ = < 1 | 0.667 | $F_{(2, 29)}$ = < 1 | 0.457 | $F_{(2, 58)}$ = 39.89 | <0.0001 | mCh<br>1st vs 4th =>0.999<br>1st vs 5th =0.006<br>4th vs 5th =0.001<br>Gq<br>1st vs 4th =>0.999<br>1st vs 5th =0.001<br>4th vs 5th =<0.0001<br>Gi<br>1st vs 4th =>0.999<br>1st vs 5th =0.013<br>4th vs 5th =0.0002 |

**Supplementary Table S9. Statistical comparisons of the effects of sustained modulation of vHPC VIP+ neurons on different elements of social behavior.** Group differences were assessed using a RM two-way ANOVA.

| | Trial x Group | | Group | | Trial | | Post-hoc |
|---|---|---|---|---|---|---|---|
| | F-value | p-value | F-value | p-value | F-value | p-value | Adjusted p-value |
| **vHPC:** Time spent "proximity" | $F_{(4, 70)}$ = < 1 | 0.894 | $F_{(2, 35)}$ = < 1 | 0.742 | $F_{(1.662, 58.18)}$ = 1.484 | 0.236 | - |
| **vHPC:** Time spent "nose-to-nose" | $F_{(4, 70)}$ = < 1 | 0.85 | $F_{(2, 35)}$ = < 1 | 0.855 | $F_{(1.537, 53.81)}$ = 3.965 | 0.035 | **mCh**<br>1st vs 4th =0.907<br>1st vs 5th =0.325<br>4th vs 5th =0.297<br>**Gq**<br>1st vs 4th =0.988<br>1st vs 5th =0.133<br>4th vs 5th =0.584<br>**Gi**<br>1st vs 4th =0.581<br>1st vs 5th =0.046<br>4th vs 5th =0.741 |
| **vHPC:** Time spent "following" | $F_{(4, 70)}$ = < 1 | 0.742 | $F_{(2, 35)}$ = < 1 | 0.76 | $F_{(1.750, 61.24)}$ = 17.48 | <0.0001 | **mCh**<br>1st vs 4th =0.978<br>1st vs 5th =0.083<br>4th vs 5th =0.087<br>**Gq**<br>1st vs 4th =0.286<br>1st vs 5th =0.218<br>4th vs 5th =0.02<br>**Gi**<br>1st vs 4th =0.138<br>1st vs 5th =0.258<br>4th vs 5th =0.002 |
| **vHPC:** Time spent "nose-to-anogenital" | $F_{(4, 70)}$ = < 1 | 0.616 | $F_{(2, 35)}$ = < 1 | 0.672 | $F_{(1.503, 52.62)}$ = 47.92 | <0.0001 | **mCh**<br>1st vs 4th =0.24<br>1st vs 5th =0.008<br>4th vs 5th =0.001<br>**Gq**<br>1st vs 4th =0.491<br>1st vs 5th =0.012<br>4th vs 5th =0.001<br>**Gi**<br>1st vs 4th =0.015<br>1st vs 5th =0.040<br>4th vs 5th =<0.0001 |

**Supplementary Table S10. Statistical comparisons of the effects of sustained modulation of VIP$^+$ neurons across corticolimbic regions on fear conditioning.** Group differences were assessed using a RM two-way ANOVA.

| | Trial x Group | | Group | | Trial | | Post-hoc |
|---|---|---|---|---|---|---|---|
| | F-value | p-value | F-value | p-value | F-value | p-value | Adjusted p-value |
| **BLA:** freezing at fear conditioning | F (8, 116) = 1.32 | 0.242 | F (2, 29) = 2.41 | 0.108 | F (2.744, 79.57) = 81.11 | <0.0001 | |
| **BLA:** freezing at test | F (2, 28) = 1.45 | 0.252 | F (2, 28) < 1 | 0.881 | F (1, 28) = 7.524 | 0.012 | - |
| **mPFC:** freezing at fear conditioning | F (8, 120) = 1.339 | 0.231 | F (2, 30) = 2.28 | 0.12 | F (2.995, 89.86) = 55.40 | <0.0001 | - |
| **mPFC:** freezing at test | F (2, 30) = 1.9 | 0.168 | F (2, 30) = 5 | 0.013 | F (1, 30) = 3.51 | 0.071 | 1st<br>mCherry vs Gi=0.932<br>mCherry vs Gq=0.018<br>Gq vs Gi=0.007<br>5th<br>mCherry vs Gi=0.2727<br>mCherry vs Gq=0.7999<br>Gq vs Gi=0.1030 |
| **vHPC:** freezing at fear conditioning | F (8, 144) = 1.59 | 0.132 | F (2, 36) = 0.0163 | 0.984 | F (2.503, 90.09) = 38.06 | <0.0001 | - |
| **vHPC:** freezing at test | F (2, 36) = 0.028 | 0.972 | F (2, 36) = 0.13 | 0.879 | F (1, 36) = 0.795 | 0.379 | - |

**Supplementary Table S11. Statistical comparisons of the effects of sustained modulation of VIP$^+$ neurons across corticolimbic regions on fear conditioning** Group differences were assessed using Kruskal-Wallis test (non-parametric data).

| | **Distribution** | **p-value** | **Significance** |
|---|---|---|---|
| **BLA:** Baseline freezing at fear conditioning | Not normal | 0.587 | ns |
| **BLA:** Baseline freezing at test | Not normal | 0.881 | ns |
| **mPFC:** Baseline freezing at fear conditioning | Not normal | 0.096 | ns |
| **mPFC:** Baseline freezing at test | Not normal | 0.001 | ** |
| **vHPC:** Baseline freezing at fear conditioning | Not normal | 0.537 | ns |
| **vHPC:** Baseline freezing at test | Not normal | 0.313 | ns |
